# Chronic stress exacerbates acute stress-induced neuronal activation in the anterior cingulate cortex and ventral hippocampus that correlates with behavioral deficits in mice

**DOI:** 10.1101/2020.02.19.956672

**Authors:** Corey Fee, Thomas Prevot, Keith Misquitta, Mounira Banasr, Etienne Sibille

## Abstract

Altered activity of corticolimbic brain regions is a hallmark of stress-related illnesses, including mood disorders, neurodegenerative diseases, and substance abuse disorders. Acute stress adaptively recruits brain region-specific functions for coping, while sustained activation under chronic stress may overwhelm feedback mechanisms and lead to pathological cellular and behavioral responses. The neural mechanisms underlying dysregulated stress response and how they contribute to behavioral deficits are poorly characterized. Here, we tested whether prior exposure to chronic restraint stress (CRS) or unpredictable chronic mild stress (UCMS) in mice could alter neuronal response to acute stress and whether these changes are associated with chronic stress-induced behavioral deficits. More specifically, we assessed neuronal activation indexed by c-Fos+ cell counts in 24 stress- and mood-related brain regions, and determined if changes in acute stress-induced neuronal activation were linked to chronic stress-induced behavioral impairments. Results indicated that CRS and UCMS led to convergent physiological and anxiety-like deficits, whereas cognition was impaired only in UCMS mice. CRS and UCMS exposure exacerbated neuronal activation in response to an acute stressor in anterior cingulate cortex (ACC) area 24b and ventral hippocampal (vHPC) CA1, CA3, and subiculum. In dysregulated brain regions, levels of neuronal activation were positively correlated with principal components capturing variance across widespread behavioral alterations relevant to stress-related disorders. Our data supports an association between a dysregulated stress response, altered corticolimbic excitation/inhibition balance, and the expression of maladaptive behaviors.

**Highlights:**

- Chronic stress models produce variable profiles of physiological deficits, anxiety-like behavior, and impaired cognition
- Acute stress-induced activation of ACC A24b & vHPC is exacerbated by prior chronic stress exposure
- In regions dysregulated by chronic stress, altered neuronal activation is positively correlated with behavioral deficits

## Introduction

The acute stress response is an adaptive mechanism that maintains homeostatic function by recruiting coordinated neuroendocrine, autonomic, and immune system activities^1^. In the brain, acute stress activates the hypothalamic-pituitary-adrenocortical (HPA) axis, releasing glucocorticoids (GCs) to mobilize energy and direct brain systems for effective coping^2^. These mediators modulate both excitatory glutamatergic pyramidal neurons and inhibitory γ-aminobutyric acid (GABA)ergic interneurons to drive changes in the functional activity of corticolimbic brain regions^3–9^. Initially adaptive, sustained activation of this response during chronic stress can overwhelm feedback mechanisms and induce neuroplastic structural and functional changes that may underlie disrupted mood and cognitive functions^10,11^. Indeed, exposure to prolonged stressors is a major precipitating factor for psychiatric disorders, including major depressive disorder (MDD) and anxiety disorders^12,13^, neurodegenerative disorders such as Alzheimer’s and Parkinson’s diseases^14^, and substance abuse disorders^15^. Further, regional activity alterations are associated with specific symptoms of stress-related disorders, including disturbed executive function in MDD^16^ and enhanced fear responses in anxiety disorders^17^. However, it remains poorly understood how dysregulation of the adaptive stress response translates to altered neuronal activity and subsequently, how regional changes in neuronal activation contribute to the development of maladaptive behaviors.

Preclinical models using exposure to chronic stress are useful tools for investigating pathological mechanisms relevant to human disorders. Among these, chronic restraint stress (CRS) and unpredictable chronic mild stress (UCMS) models are most common due to their construct, predictive, and face validity for modeling cell- and circuit-level pathological changes together with behavioral alterations related to human symptoms^18–21^, that are reverse by pharmacological treatments. For example, rodents exposed to chronic stress display neuronal atrophy in the prefrontal cortex (PFC) and hippocampus (HPC), but hypertrophy in the amygdala, concurrent with anxiety- and depressive-like behavioral deficits^22–29^, and closely resembling findings in humans with mood disorders^30–32^. That synaptic changes are altered in a brain region-specific manner implies a putative imbalance in neuronal activity and function. Indeed, chronic stress negatively impacts both glutamatergic and GABAergic neurons (reviewed in^33^), supporting theories that sustained activation leads to altered excitation/inhibition balance (EIB) in cell circuits. These changes are also bi-directional. For instance, PFC and HPC feedback mechanisms regulate initiation and termination of the HPA stress response^34,35^. Therefore, functional deficits emerging from altered activity of corticolimbic brain regions likely represent both a cause and consequence of sustained challenge on the adaptive stress response^36^.

Neuroimaging studies validate structural and functional changes in corticolimbic brain regions of patients with stress-related disorders^37,38^. However, deriving insights about chronic stress is less straightforward due to variability introduced by patient characteristics (e.g., medication history, comorbid disorders, timing and severity of stress exposure) and experimental conditions (e.g., resting state vs. task-related activity). Thus, rodent studies using systematic activity mapping after acute and chronic stress have been useful to understand functional consequences of stress-related pathology. For example, functional magnetic resonance imaging (fMRI) studies in CRS mice identified dendritic remodeling of the PFC, HPC, and amygdala with altered functional connectivity across regions of the default mode network^39^, in which activity changes are linked to symptom emergence in MDD^40^, PTSD^41^, Parkinson’s disease^42^, and substance abuse disorders^43^. Immediate early gene (IEG) markers, such as c-Fos, similarly yield high spatiotemporal resolution for characterizing functional alterations induced by chronic stress. Stress-induced recruitment of brain region activity initiates c-Fos expression, which varies depending on the type of stress, severity, and duration of exposure^44,45^. Additionally, patterns of acute activation can become modulated by prior exposure to chronic stress, potentially reflecting functional disturbances. However, findings are mixed as some studies report sensitization of the acute stress response by prior chronic stress exposure, reflected by enhanced c-Fos levels in the anterior cingulate cortex (ACC), HPC, hypothalamus, and amygdala^46,47^, while others report response habituation and c-Fos attenuation following acute stress in distinct and overlapping regions^48–50^.

Here, we employed two mouse models of chronic stress exposure, longitudinal and end-point behavioral phenotyping, and comprehensive mapping of regional activation via c-Fos protein immunohistochemistry (IHC) following an acute stressor subsequent to chronic stress exposure. We sought to determine (a) whether chronic stress induced behavioral alterations along dimensions including physiological deficits, anxiety- and anhedonia-like behaviors, antidepressant-predictive behaviors, and cognitive impairments (b) whether region-specific responses to acute stress were dysregulated by prior chronic stress exposure and (c) whether regional activation changes were associated with behavioral deficits relevant to stress-related disorders. Control, CRS, and UCMS mice were used to avoid model-specific conclusions about the effects of chronic stress on behavioral and functional outcomes^44,45,51^. Behavioral phenotyping assessed deficits along symptom-related dimensions, including physiological alterations, anxiety- and anhedonia-like behaviors, antidepressant-predictive behaviors, and cognitive impairments. Semi-quantitative c-Fos mapping was performed via IHC in 24 stress- and mood-related brain regions (plus motor cortex and white matter control regions) identified via *a priori* literature review^37,52–55^. Contributing relationships were investigated via correlational analyses between regional neuronal activation and unbiased parameters that captured variance across behavioral deficit dimensions. We hypothesized that prior chronic stress exposure alters acute stress-induced neuronal activation in corticolimbic brain regions, and that these changes correlate with behavioral deficits. Our findings revealed that CRS and UCMS converged to exacerbate levels of neuronal activation by acute stress in ACC and vHPC subregions. These changes were positively correlated with stress-related behavioral dimensions, revealing a pathway by which the acute stress response becomes dysregulated and contributes to the expression of behavioral deficits.

## Experimental Procedures

### Animals

Eight-week C57BL/6J mice (Jackson Laboratories, Bar Harbor, ME) were kept under a 12/12h light-dark cycle with food and water *ad libitum*. Group-housed mice underwent 2-week facility habituation before random assignment to control (daily handling), CRS, or UCMS groups (*n*=12-14/group; 50% male/female). CRS/UCMS mice were single-housed at chronic stress initiation, whereas control mice were single-housed prior to end-point testing. All tests were performed in accordance with Canadian Council on Animal Care (CCAC) guidelines.

### Chronic Stress Procedures

UCMS mice received 5 weeks of randomized mild stressors (2-4/day), and 3 weeks maintenance (1-2/day) during testing, based on validated methods^56^. Stressors included: forced bath, wet bedding, predator odor, light cycle reversal/disruption, cage switching, tilted cage, reduced space, acute restraint, and enrichment removal. CRS mice received 5 weeks acute restraint (1h; 2x/day), and 3 weeks maintenance (1x/day) during testing, based on validated methods^57^.

### Animal testing

Mice were tested in randomized order by experimenters blind to group assignments. Tests were performed during the light phase on non-consecutive days and measured *physiological changes* i.e., coat state, weight gain, *anxiety-like behavior* i.e., PhenoTyper Test (PT), Elevated Plus-maze (EPM), Open Field Test (OFT), and Novelty-Suppressed Feeding (NSF), *anhedonia-like behavior* i.e., Sucrose Consumption Test (SCT), or *both* i.e., Novelty-Induced Hypophagia test (NIH) (**Supplementary Fig. 1**). Mice were also assessed on the *antidepressant-predictive* Forced Swim Test (FST), and for *cognitive impairment* i.e., Y-maze (YM) and Novel Object Recognition Test (NORT). To characterize the induction of stress-related deficits, *Longitudinal tests* (i.e., repeatable assessments not relying on novelty^58^) for weight, coat state, PT, and SCT were administered weekly, 6x from week 0 (before stress) to week 5 (5^th^ week of chronic stress). To characterize the full extent of chronic stress-induced deficits, *End-point* tests included all tests mentioned previously, performed on non-consecutive days from weeks 6-8 (after 5+ weeks chronic stress).

### Physiological tests

Animals’ coat state quality was scored across 7 anatomical areas 0, 0.5, or 1, from maintained to unkempt, and a sum was calculated^59^. Weight was tracked weekly and expressed as percent change (%) to measure deviation from normal development. LA was assessed for 25min in a home cage-like environment with an overhead camera and tracked offline using ANY-maze software (Stoelting Co., Wood Dale, IL).

### Anxiety-like behavior tests

Tests included the PT, a test developed by our lab to assess anxiogenic response to a light challenge in a home cage-like environment^58^. Time spent in a PhenoTyper apparatus (Noldus, Leesburg, VA), with designated food (6.5×15cm) and shelter (10×10cm) zones was tracked during the dark cycle (7pm-7am). At 11pm, an aversive spotlight was applied over the food zone for 1h. All mice spend more time hiding in the shelter during this challenge, however, chronic stress-exposed mice show a protracted return to normal exploration after the challenge ends, reflected by increased shelter zone time and decreased food zone time for up to 10h^58^. Anxiogenic shelter/food zone response was represented by an area under the curve (AUC): the summed average of 1h responses from the start of light challenge to end of the test (~8h). In the EPM, mice were recorded for 10min exploring an elevated (55cm) cross-shaped maze with two open arms (27×5cm) and two partially-enclosed arms (27×5×15cm), in a dimly lit room. ANY-maze software was used to measure time spent (s) and percentage of distance travelled (%, cm) in open and closed arms, excluding the neutral middle zone. In the OFT, mice were recorded for 10min exploring an open arena (70×70×33cm) in a dimly lit room. ANY-maze software delineated 3 concentric squares of equal area and tracked time spent (s) in the innermost center zone (40×40cm) and outermost (70×70cm perimeter zone. In the NSF, food-deprived (16h) mice were timed for latency to approach and feed on a food pellet (s) in a brightly lit arena (62×31×48cm). Afterwards, home cage approach and feed latency was tracked to assess potential bias due to appetitive drive^60^.

### Anhedonia-like behavior tests

Tests included NIH, involving two days reward habituation (1mL of sweetened condensed milk), followed by measuring home cage reward approach latency on the third day as a measure of reward drive. On the fourth day, reward approach latency was measured in a novel brightly-lit cage, by an experimenter. The novel environment contributes to NIH being considered as an index of both anxiety- and anhedonia-like behaviors^61^. In the SCT, home cage water bottles were replaced with sucrose solution (1%; Sigma, St. Louis, MO) for 48h. Mice were then fluid-deprived overnight (~14h), and sucrose intake measured for 1h. 2 days later, this process was repeated for water. Results were interpreted as a sucrose preference percentage (sucrose consumed/sucrose+water consumed, %), to control for potential bias due to fluid intake.

### Antidepressant-predictive behavior tests

Tests included the FST, a swim test that assesses chronic stress-induced increased immobility that is reversed by antidepressant treatment^62^. Mice underwent a 6min swim trial in a glass beaker (25×16cm) of room temperature water and were then dried and warmed by a heat lamp. Trials were recorded and assessed offline for total immobility time, scored manually by an experimenter, blind to group assignments.

### Cognitive tests

Tests included the YM, a black plastic Y-maze with 3 arms (26×8×13cm) separated by 120^°^ and gated by sliding doors. YM leverages the exploratory nature of rodents to alternate between 2 goal arms, across 7 successive trials^63^. Mice were habituated to the apparatus and distal cues for 10min sessions on 2 consecutive days. On day 3, mice underwent 7-trials to assess goal arm alternation with a non-challenging 30s inter-trial interval (ITI). On day 4, alternation was assessed across 7-trials with a 90s ITI, simulating increased cognitive load. An 8^th^ trial with 5s ITI was added to assess loss of motivation; mice failing this were excluded. Correct alternation percentage reflected an index of working memory performance (50% = random). The NORT leverages rodent’s preference for novelty to assess short-term memory (recognition/recall)^64^. Mice were habituated to an empty open field (70×70×33cm) and then to the field with 2 identical plastic objects (“familiar”) over 10min trials on consecutive days. On day 3, mice underwent a familiar pre-trial: tracking time spent with 2 identical familiar objects and then 3h later were tested for recall in the trial: switching one familiar object for a novel object. A discrimination ratio measured time with left familiar object/time with both (pre-trial) or time with novel object/time with both (trial).

### Immunohistochemistry

Forty-eight hours after end-point testing, mice received an acute stressor (5min 18^°^C cold swim) to stimulate c-Fos expression. Ninety minutes later, mice were deeply anesthetized with avertin (125mg/kg, i.p.) and transcardially perfused with 50mL 0.1M phosphate-buffered saline (PBS, pH 7.4), followed by 75mL 4% paraformaldehyde (PFA) in PBS. Extracted brains were post-fixed in PFA for 24h at 4^°^C, cryoprotected in 30% sucrose in PBS for 2 days, then frozen on dry ice and cryosectioned at 40μm. Ten sets of full brain sections were collected yielding ~15 sections/set/mouse processed for IHC.

Neuronal activation was quantified via c-Fos protein immunolabeling ^65^. Free-floating sections were rinsed in PBS, pretreated with 3% H_2_O_2_ for 20min, and permeabilized in 0.3% Triton X-100 PBS for 20min. Sections were again rinsed in 0.3% Triton-PBS before applying 10% normal goat serum (NGS) in 0.3% Triton-PBS as blocking agent for 30min and incubated in primary antibody (rabbit anti-c-Fos 226003, 1:200 in 3% NGS/0.3% PBS-Triton; Synaptic Systems, Germany) for 72h at 4^°^C. Sections were rinsed with PBS and incubated with secondary antibody (biotinylated goat anti-rabbit IgG BA-1000, 1:500 in 0.3% Triton-PBS; Vector Laboratories, Burlingame, CA) for 2h at room temperature. Reactions were enhanced with Vectastain ABC/HRP kit (Vector Laboratories) for 2h, then rinsed and detected with 0.025% 3,3’-diaminobenzidine (DAB) and 0.025% of H_2_O_2_ in 0.5M Tris-buffered saline (pH 7.4). After rinsing, sections were mounted, dehydrated, and coverslipped using DEPEX medium.

### Microscopy

Quantitative analysis of c-Fos+ cell counts was performed using ZEN 2 software on scans acquired with a Zeiss AxioScanZ1 slide scanner (ZEISS Microscopy, Oberkochen, Germany). Regions were delineated at low magnification (2X) according to the Franklin and Paxinos Mouse Brain Atlas^66^. Cell counts were acquired in a semi-automated manner across full resolution 20X images using ZEN 2 with consistent thresholds for nuclei size, roundness, signal intensity, and segmentation. Cell counts for each region were obtained bilaterally across 2-3 evenly spaced (~400μm) sections and expressed as mean cell density count (per 10,000 μm^2^) per mouse per region, based on previous studies^67^. As a summary measure, regional cell density counts were produced by averaging across subregions, where >2 subregions were quantified (i.e., PFC, amygdala, dHPC and vHPC, and brainwide).

### Statistical Analysis

Data are expressed as mean ± standard error of the mean (SEM). Data were analyzed using SPSS (IBM, Armonk, NY). Behavioral parameters and cell counts were analyzed using one-way analysis of covariance (ANCOVA) with group as independent variable and sex as covariate, followed by Scheffe *post hoc* analysis. Time-course parameters from the coat state, weight gain, SCT, PT, and LA assessments were assessed by repeated measures ANCOVA for within- and between-subject comparisons using Greenhouse-Geisser correction for parameters failing sphericity assumptions. To address variability associated with classical rodent behavior tests^68,69^, we employed principal components analysis (PCA) to generate summary scores reflecting overall depressive-like behavioral deficits, based on previous methods^56,58^. Twenty-six parameters measuring baseline (i.e., week 0 or week 6-9 pre-trial readouts) and end-point (i.e., weeks 6-8 trial readouts) behaviors were Z-normalized^68^ and loaded into PCA with varimax rotation, regarding mice as one experimental unit. For interpretation, parameter loadings onto principal components (PCs) were examined, wherein those > 0.3 (absolute value) were considered significant based-on critical *r* values corresponding to *p* < .05 for the current sample size. Pearson’s *r* was used to assess relationships between cell density and behavioral PCs.

## Results

### Repeatable tests revealed progressive physiological and behavioral deficits during 5 weeks of chronic stress

Mice were assessed weekly for weight gain, fur coat deterioration, SCT, and PT from weeks 0 to 5, including 1-week baseline and 5-weeks chronic stress (**Fig. 1**). Repeated measures ANCOVA of weight gain revealed a significant main effect of group (*F_2,36_* = 19.92;*p* < .001), time (*F_3.8,134.19_* = 3.99;*p* < .01), and group*time interaction (*F_7.6,332.73_* =4.95; *p* < .001) (**Fig. 1A**). *Post hoc* analysis revealed that CRS altered weight gain relative to controls (*p* < .01), with significantly decreased change in weight gain observed the 1^st^, 4^th^, and 5^th^ week of stress. ANCOVA revealed significant sex differences (*p* <. 01) wherein females gained less weight across groups (**Supplementary Table 1**). Analysis of coat state scores showed a significant main effect of group (*F_2,36_* = 16.34;*p* < .001), time (*F_2.94,105.83_* = 78.96;*p* < .001), and a group*time interaction (*F_5.88,105.83_* = 9.49; *p* < .001) (**Fig. 1B**). *Post hoc* analysis showed that CRS and UCMS exposure increased coat state deterioration from weeks 1 to 5 (*ps* < .01). ANCOVA revealed significant sex differences, wherein deterioration was more extensive in males (*p* < .001; **Supplementary Table 1**).

**Fig 1.**
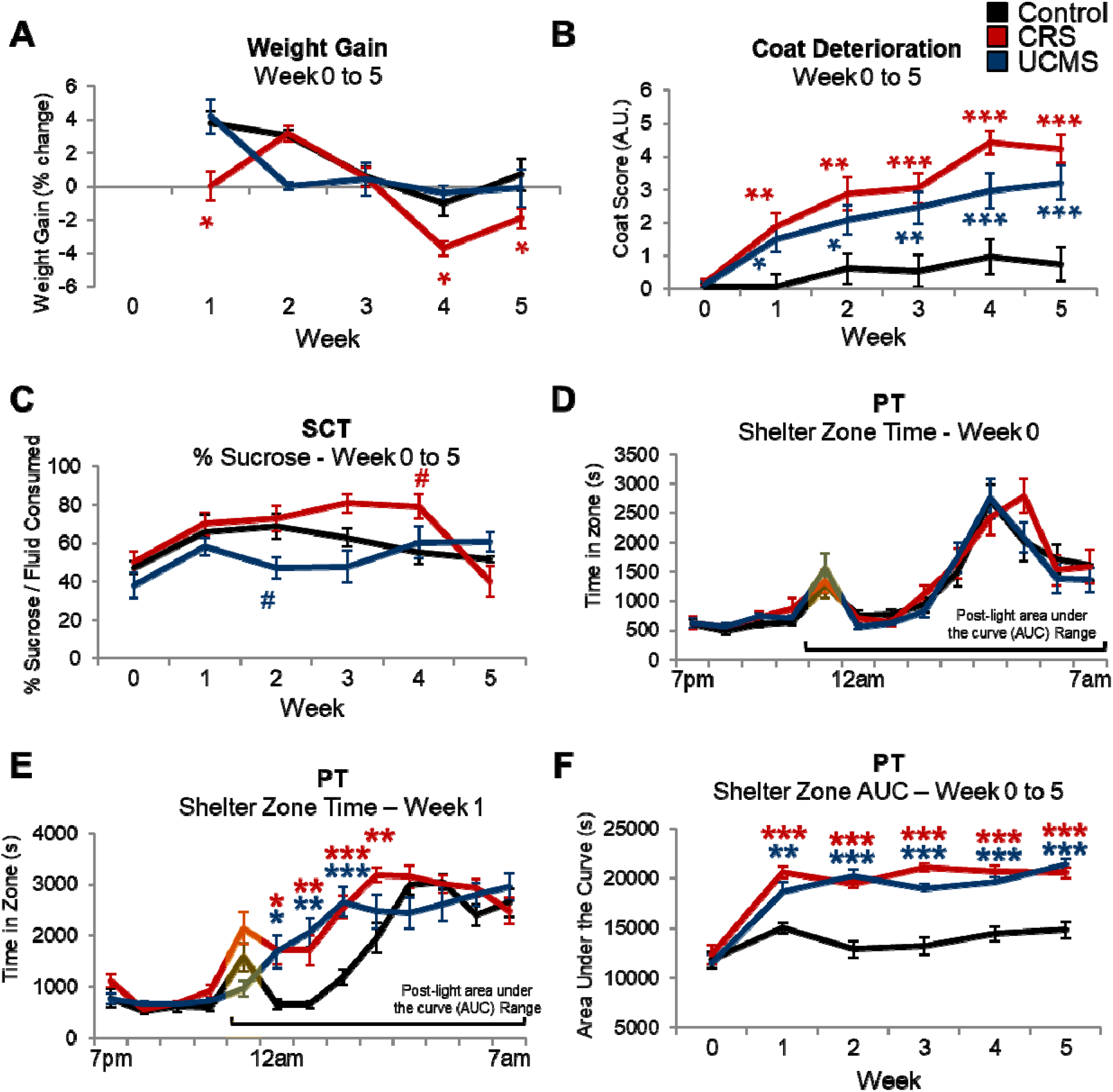
Longitudinal analysis using weekly repeatable physiological and behavior tests to confirm induction of stress-related deficits in mice assigned to control (daily handling), CRS, or UCMS conditions (*n*=12-14/group; 50% male/female). (A) Percentage weight gain. (B) Fur coat state deterioration scores (AU = arbitrary units). (C) Percent sucrose of total volume consumed in the sucrose consumption test (SCT). (D) Time spent in the shelter zone of the PhenoTyper Test (PT) in week 0 prior to chronic stress. (E) Time spent in the shelter zone of the PT after 1 week of chronic stress. (F) Area under the curve (AUC) for shelter zone time in PT from the beginning of the light challenge (yellow vertical bars) until the end of the test. *** *p* < .001; ** *p* < .01; \**p* < .05; ^#^*p* < .1 for CRS (red) or UCMS (blue) vs. controls.

Analyses of week 0 to 5 sucrose preference ratio in the SCT revealed a significant main effect of group (*F_2,34_* = 10.02;*p* < .001), time (*F_5,175_* = 4.98;*p* < .001), and group*time interaction (*F_10,175_* = 2.34;*p* < .05) (**Fig. 1C**). *Post hoc* comparisons revealed trend-level differences for CRS and UCMS mice vs. control mice (*ps* = .08) and a significant decrease in CRS vs. UCMS mice (*p* < .001). However, only trend-level differences were found at individual time points, including decreased sucrose preference for UCMS vs. CRS and control mice week 2 (*ps =* 0.6-0.7), and increased sucrose preference for CRS vs. control mice week 3 (*p* = .052). No main effect or interaction with sex was found.

PT was used to assess shelter zone activity before, during, and after a 1h light challenge from week 0 to 5 (**Fig. 1D-F**). In week 0 (prior to stress), repeated measures ANCOVA revealed a significant effect of time (*F_4.81,173_* = 48.09;*p* < .001) on shelter zone time, but no effect of group or their interaction (*ps* > .4) (**Fig. 1D**). In week 1, main effects for time (*F_6.63,228.86_* = 59.7; *p* < .001), group (*F_2,36_* = 22.49 *p* < .001), and group*time interaction (*F_12.771,228.86_* = 3.92;*p* < .001) were found for shelter zone time (**Fig. 1E**). *Post hoc* analysis revealed significantly increased shelter zone time for CRS and UCMS mice relative to controls for each of the 4 hours following the light challenge (*ps* < .05). Shelter Zone AUCs were calculated to summarize anxiogenic light responses from week 0 to 5 PTs (**Fig. 1F**). Repeated measures ANCOVA of AUC data revealed significant main effects of time (*F_2.65,95.39_* = 57.09;*p* < .001), group (*F_2,36_* = 43.11;*p* < .001), and group*time interaction (*F_5.3,228.86_* = 7.65;*p* < .001). Relative to control mice, CRS and UCMS mice spent more time in the shelter zone following the light challenge from weeks 1 to 5 (*ps* < .01). No main effect or interaction with sex was found.

### End-point tests revealed induction of physiological deficits and anxiety-like behavior in CRS-exposed mice and physiological deficits, anxiety-like behavior, and cognitive impairment in UCMS-exposed mice

End-point physiological measurements included final assessments of coat state and weight gain, plus LA (**Fig. 2A-C**). One-way ANCOVA revealed significant group effects on coat state (*F_2,33_* = 61.67; *p* < .001) and weight gain (*F_2,33_* = 6.07;*p* < .01). CRS and UCMS significantly induced coat state deterioration (*p* < .001) (**Fig. 2A**) in males and females. A sex (*F_1,33_* = 6.07;*p* < .001) and sex*group interaction (*F_2,33_* = 2.95;*p* < .05) was found wherein females overall and CRS/UCMS-exposed females relative to males had less deterioration (**Supplementary Table 1**). For % weight gain, significant effects of group (*F_2,33_* = 28.8;*p* < .001), sex (*F_1,33_* = 103.53;*p* < .001), and group*sex (*F_2,33_* = 4.36;*p* < .05) were found showing that, relative to controls, weight gain was less extensive in CRS males and females (*p* < .001), but only UCMS males (*p* < .001) (**Fig. 2B**; sex effects **Supplementary Table 1**). Total distance travelled in the LA test was not significantly affected by group, sex, or their interaction (**Fig. 2C**).

**Fig 2.**
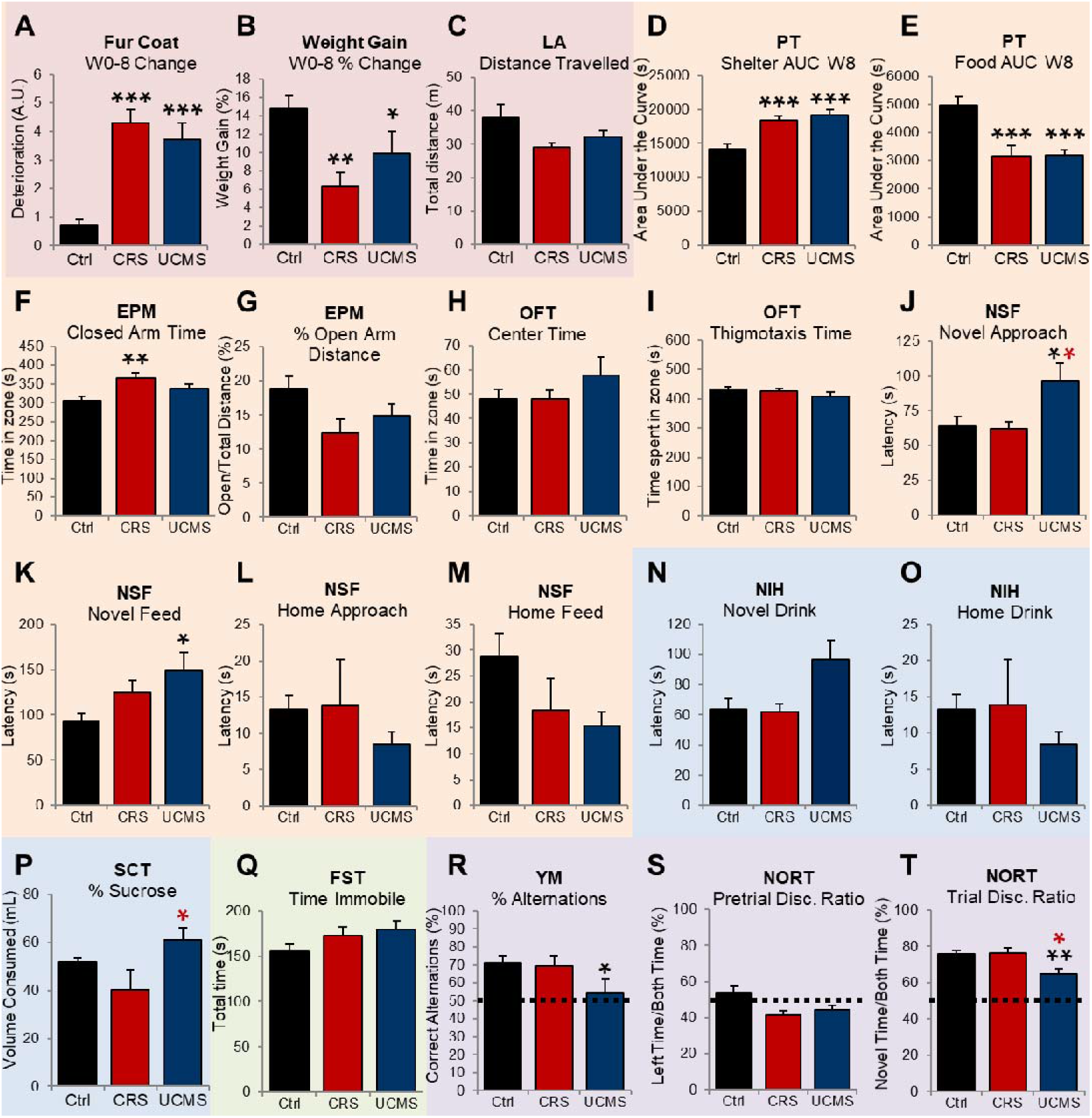
End-point analysis using multiple behavior tests to characterize induction of deficits along physiological (light red), anxiety-like (light orange), anhedonia-like (blue), antidepressant-predictive (light green), and cognitive impairment (purple) dimensions in control, CRS, and UCMS mice (*n*=12-14/group; 50% male/female). (A) Fur coat change from week 0 to 8. (B) Percent weight gain from week 0 to 8. (C) Distance travelled in 25 min locomotor activity (LA) test. (D) Area under the curve (AUC) of shelter zone time. (E) Shelter zone and (F) food zone time area under the curve (AUC) in PhenoTyper Test (PT) from the beginning of the light challenge until the end of the test. (F) Time spent in closed arms and (G) % of distance travelled in open arms in elevated-plus maze (EPM). (H) Time spent in the center and (I) thigmotaxis zones in open-field test (OFT). (J) Time to approach and (K) feed on a food pellet in novel environment, and (L) approach and (K) feed on a pellet in homecage, in the novelty-suppressed feeding test (NSF). (N) Latency to drink in a novel and (O) homecage environment in the novelty-induced hypophagia test (NIH). (P) Percent sucrose of total volume consumed in the sucrose consumption test (SCT). (Q) Total time immobile in the forced-swim test (FST). (R) Percent correct alternations from 7-trials in the Y-maze (YM). (S) Discrimination ratio for pre-trial with identical objects and (T) trial with familiar/novel objects in the novel object recognition test (NORT). *** *p* < .001; ** *p* < .01; \**p* < .05 for indicated group vs. control (black), CRS (red) or UCMS (blue).

End-point anxiety-like behavior was assessed in the PT, EPM, OFT, and NSF. In the PT, group assignment had a significant effect on shelter (*F_2,33_* = 18.56;*p* < .001) and food zone AUCs (*F_2,33_* = 11.51; *p* < .001) (**Fig. 2D-E**). The light challenge significantly increased shelter zone time and decreased food zone time in CRS and UCMS mice relative to controls (*ps* < .001), suggesting persistent avoidance of the previously lit zone and reflecting a protracted anxiogenic response. A significant sex effect was found for shelter zone AUC (*F_1,33_* = 7.84; *p* < .01), but *post hoc* analysis revealed only a trend-level increase for males (*p* = .064; not shown). In the EPM, group assignment significantly altered time spent in closed arms (*F_2,35_* = 7.1;*p* < .01), without changes in time spent (not shown) or % of distance travelled in open arms (**Fig. 2F-G**). Closed arm time was significantly elevated in CRS mice relative to controls (*p* < .01). In the OFT, center zone and outermost zone time were not affected by chronic stress (**Fig. 2H-I**). In the NSF, we found a main effect of group on latency to approach (*F_2,36_* = 4.87;*p* < .05) and feed (*F_2,36_* = 3.66; *p* < .05) on a food pellet in the novel arena (**Fig. 2J-K**). UCMS increased latency to approach relative to CRS and control mice (*ps* < .05) and latency to feed relative to control mice only (*p* <.05). Home cage latencies were unchanged, reflecting no bias due to appetitive drive (L-M). Sex was not a significant covariate in the EPM, OFT, or NSF.

End-point anhedonia-like behavior were assessed using reward approach and consumption in NIH and SCT. In the NIH, chronic stress did not significantly affect latency to drink milk in novel or home cage tests (**Fig. 2N-O**). Sex was a significant covariate (*F_2,33_* = 5.91;*p* < .05) as females had increased latency to drink in a novel environment relative to male mice (*p* < .05; **supplementary table 1**). In the SCT, sucrose preference ratio was significantly affected by group (*F_2,36_* = 3.42;*p* < .05), but not sex. CRS mice had decreased consumption relative to UCMS mice (*p* < .05) (**Fig. 2P**).

End-point measures of working and short-term memory were assessed in the YM and NORT, respectively. In YM, one-way ANCOVA revealed significant effects of stress on alternation rate (*F_2,36_*= 4.48;*p* < .05) (**Fig. 2R**). Specifically, under cognitive load of 90s ITI, UCMS mice alternated arm choice less than controls (*p* < .05), suggesting a working memory deficit. In the NORT, pre-trial discrimination ratio (measuring time spent with 1 of 2 identical objects) was not significantly affected by stress (*p* > .12; **Fig. 2S**), indicating that no arena side preference existed. Short-term recall was assessed 3h later by trial discrimination ratio (measuring novel object/novel+familiar object time) indicating a significant group effect for object recall (*F_2,36_* = 7.13;*p* < .01). UCMS significantly decreased discrimination ratio compared to control and CRS mice (*ps* <.05) (**Fig. 2T**). Sex was not a significant covariate in the YM or NORT.

### Principal components analysis reveals 2 components associated with stress-induced variance across behavioral tests

As an alternative approach to assessing individual behaviors, we used dimension reduction via PCA to generate parameters capturing behavioral variance across tests. PCA was conducted on 26 z-normalized parameters (including baseline and end-point readouts) revealing a 5-component solution accounting for 56.7% of the total variance (**Supplementary Fig. 2**). One-way ANCOVA revealed a significant main effect of group on the top two components: PC1 (*F_2,36_* = 15.2;*p* < .001, 16.43% of variance) and PC2 (*F_2, 36_* = 3.29; *p* < .05, 14.27% of variance) (**Fig. 3A-B**), with no effect of sex or their interaction. Relative to control mice, CRS and UCMS mice had significantly higher PC1 scores (*p* < .001), whereas only CRS mice had significantly higher PC2 scores (*p* < .05). Strong loadings were found for parameters of physiological and anxiety-like deficits for both PC1 (**Fig. 3C**) and PC2 (**Fig. 3D**), indicating that these components captured emotional reactivity across behavioral dimensions and groups. However, qualitative inspection revealed that PC1 loaded stronger on physiological, antidepressant-predictive, and cognitive deficits, whereas PC2 loaded mainly on anxiety-like deficits and negatively loaded on cognitive deficits, consistent with reflecting variance solely from CRS mice (**Fig. 3A-B**). PC3-5 were independent of group and/or sex effects, reflecting variance from other unrelated sources.

**Fig 3.**
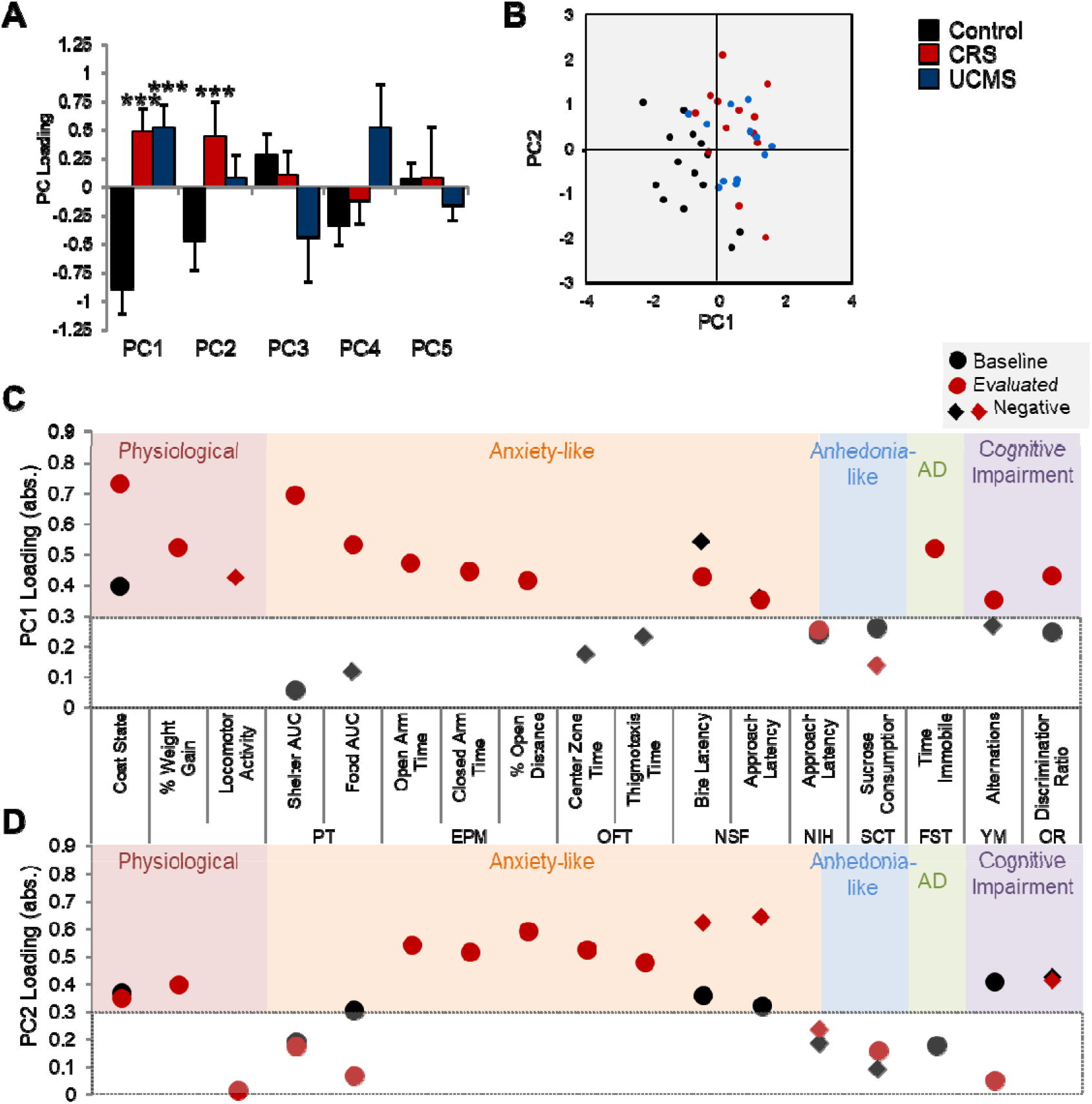
Dimension reduction via principal component (PC) analysis of behavioral data from mice at baseline (week 0 or pre-trial conditions) or evaluated (weeks 6-8 trial conditions) states. (A) PC loadings by group indicating captured behavioral variance related to chronic stress for CRS and UCMS mice for PC1 and CRS mice for PC2. Absolute values are reported; diamonds = negative loadings. (C) Visualization of clustering by groups and PC1/2. (D) PC1 and (E) PC2 loadings for individual tests demonstrating that both components captured variance across multiple symptom-like behavioral dimensions. *** *p* < .001; ** *p* < .01; * *p* < .05 for indicated group vs. controls.

### Chronic stress exacerbates the effect of acute stress on c-Fos+ cell density in prefrontal cortex and ventral hippocampal subregions

We next assessed the impact of prior chronic stress exposure on levels of c-Fos+ cell density induced by acute stress. We first established the ability for acute stress to induce c-Fos activation via analysis of c-Fos+ cell density in 26 brain regions, including 24 stress- and mood-related brain regions, 2 control regions (motor cortex and white matter), plus 5 summary measures (total PFC, Amygdala, dHPC, vHPC, and brainwide) in a separate cohort of naïve adult C57BL/6J mice following an acute swim stress or normal home cage environment (*n*=6/group; 50% male/female). Acute stress induced significant elevations in c-Fos+ cell density in 15/26 regions examined, including the PFC (areas 24a, 24b, and 32), basomedial amygdala, dorsal HPC (CA1), ventral HPC (CA1, CA3, subiculum), nucleus accumbens, lateral septum, bed nucleus of stria terminalis, lateral hypothalamus, periaqueductal gray, entorhinal cortex, and motor cortex (*ps* < .05), but not the basolateral amygdala, dorsal dentate gyrus, caudate putamen, ventromedial nucleus, ventral tegmental area, insular cortex, perirhinal cortex, or white matter (**Supplementary Fig. 3**). Acute stress induced trend-level elevations in c-Fos+ cell density in the central amygdala and the dorsal HPC (CA2, CA3) (*p* < .1).

To test whether chronic stress alters the induction of c-Fos activation, control, CRS, and UCMS mice from behavioral experiments were then subjected to the same acute swim stressor and brains were extracted and processed for IHC (**Fig. 4A**). Significant main effects of group on c-Fos+ cell density were observed only in area 24b of the PFC (*F_2,36_* = 4.11; *p* < .05), overall ventral hippocampus (vHPC; *F_2,36_*= 6.47;*p* < .01) and specifically vHPC CA1 (*F_2,36_* = 4.92;*p* < .01), CA3 (*F_2,36_* = 5.97;*p* < .01) or subiculum (vSub; *F_2,36_* = 3.75;*p* < .05) (**Fig. 4B-I**). *Post hoc* analyses revealed significant elevations in acute stress-induced neuronal activity among CRS and UCMS mice relative to controls in A24b (**Fig. 4A**), vCA1, and vCA3 (**Fig. 4D**), and in CRS mice relative to controls in vSub and overall vHPC (*ps* < .05). No main effect or interaction with sex was found.

**Fig 4.**
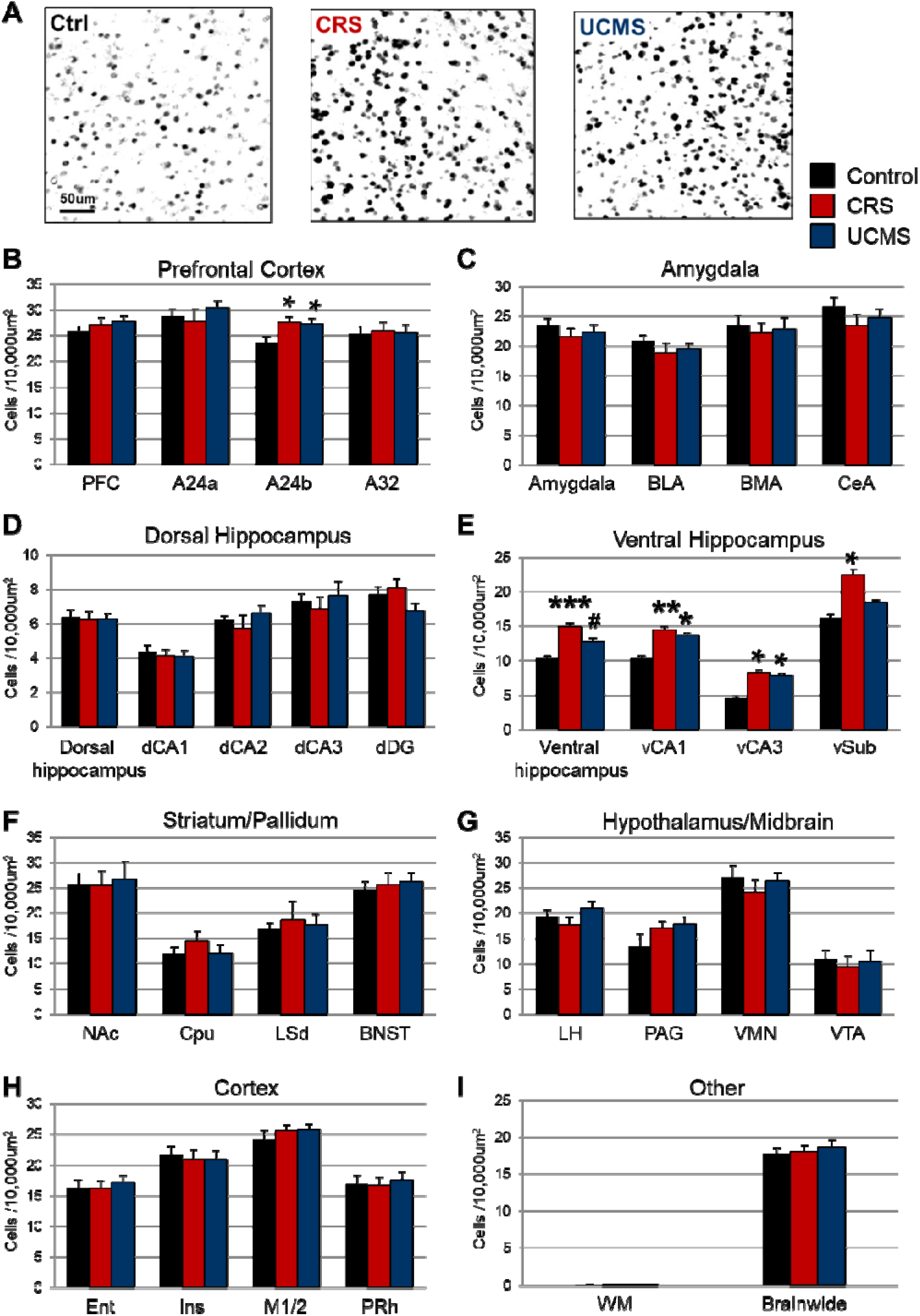
Neuronal activation quantified regionally via absolute density counts for c-Fos+ cells 90 min after exposure to an acute stressor in control, CRS, and UCMS mice (*n*=12-14/group; 50% male/female). (A) Representative qualitative images of c-Fos staining in A24b (8-bit, thresholded). (B) prefrontal cortex (PFC) as an average index of A24a, A24b, and A32. (C) Amygdala as an average index of basolateral (BLA), basomedial (BMA), and central (CeA) parts. (D) Dorsal hippocampus (dHPC) as an average index of CA1, CA2, CA3, and dentate gyrus (DG). (E) Ventral hippocampus (vHPC) as an average index of CA1, CA3, and subiculum (sub). (F) Striatum/pallidum groups comprising nucleus accumbens (NAc), caudate putamen (Cpu), lateral septum (LSd), and bed nucleus of stria terminalis (BNST). (G) Hypothalamus/midbrain groups comprising lateral hypothalamus (LH), periaqueductal gray (PAG), ventromedial nucleus (VMN), and ventral tegmental area (VTA). (H) Other cortical regions including entorhinal (Ent), insular (Ins), motor (M1/2), and perirhinal (PRh) cortex. (I) Control and overall regions: white matter (WM) and brain-wide (average of all regions).. \*\*\**p* < .001; ** *p* < .01; \**p* < .05 for indicated group vs. controls.

These results demonstrate that neuronal activation induced by an acute swim stressor is exacerbated by prior chronic stress exposure in the PFC and vHPC, when compared to acute swim stress-induced neuronal activation in all other tested brain regions tested.

### Levels of neuronal activation in prefrontal A24b and vHPC subregions positively correlate with stress-related behavioral components

We next sought to investigate associations between behavioral component and neuronal activation via correlational analyses of cross-dimensional principal components and z-normalized c-Fos**+** cell densities in subregions affected by chronic stress (**Fig. 5A-B**). These analyses revealed significant positive correlations between PC1 and c-Fos levels in A24b (*r* = .329, *p* = .041) and vCA3 (*r* = .506, *p* = .007) (**Fig. 5A**), as well as PC2 and c-Fos levels in A24b (*r* = .361,*p* = .024), vCA1 (*r* = .588,*p* < .001), and vSub (*r* = .447, *p* = .032) (**Fig. 5B**). For all comparisons, mice with higher levels of regional c-Fos activation had greater PC1 or PC2 scores, reflecting increased behavioral emotionality. Sex was not a significant covariate in these analyses.

**Fig 5.**
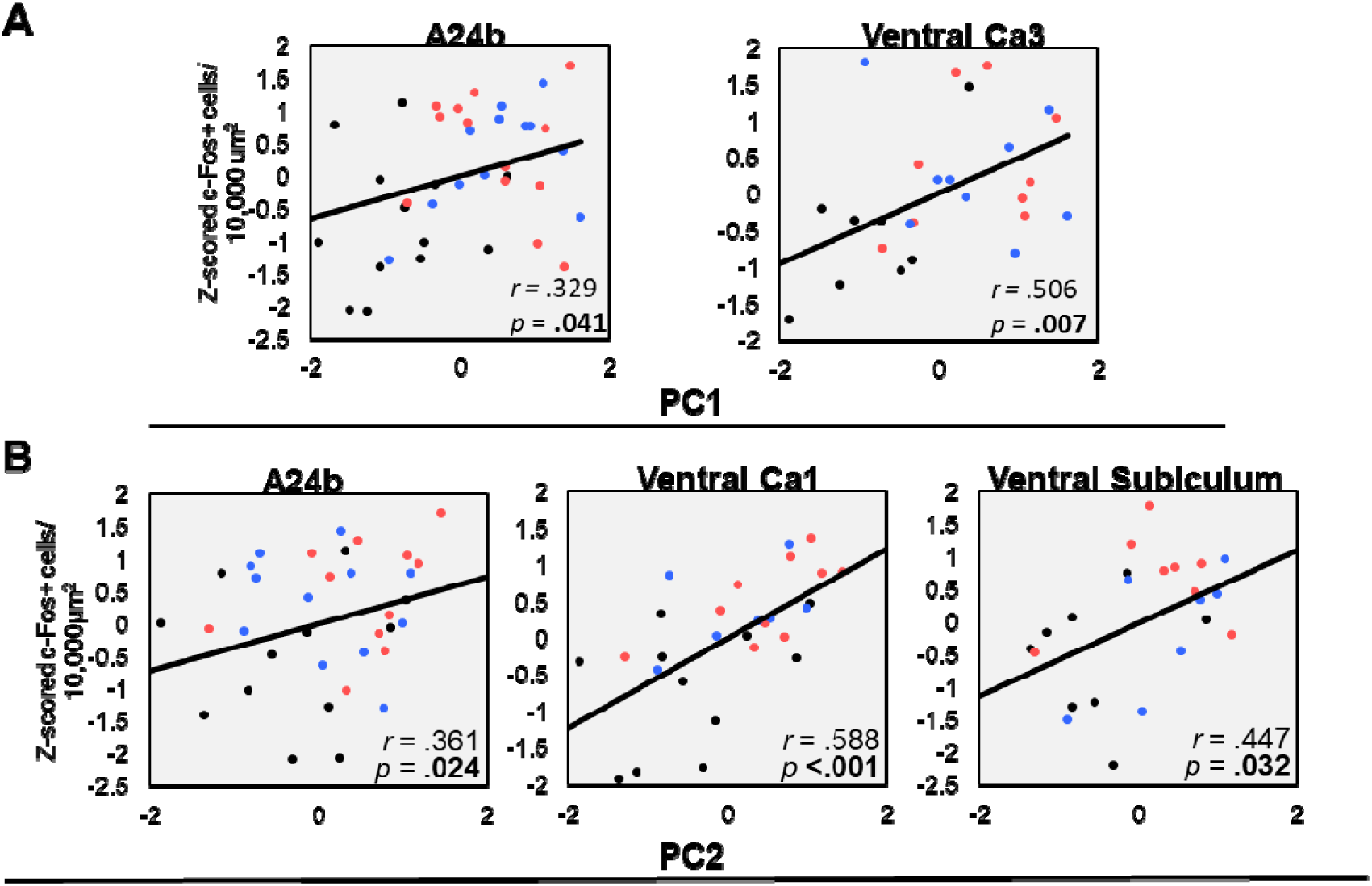
Correlational analysis of acute stress-mediated c-Fos+ cell density (z-scored) in regions dysregulated by chronic stress (A24b, vCA1, vCA3, vSub) and principal components (PC) of trans-dimensional stress-related behavioral alterations. (A) PC1 and (B) PC2 correlations with c-Fos+ cell density among control (black), CRS (red), and UCMS (blue) mice. Associations between PC1 and vCA1/vSub or PC2 and vCA3 were not significant (not shown).

## Discussion

In this report, we characterized the effect of prior chronic stress exposure on brain region-specific neuronal activation induced by acute stress and investigated associations with behavioral deficits relevant to symptoms of human stress-related disorders. We first demonstrated that two prominent chronic stress models, CRS and UCMS, lead to convergent progressive outcomes over 8 weeks of exposure in male and female mice. This included physiological and anxiety-like deficits in both groups, and cognitive impairment only in UCMS mice. To capture common elements underlying behavioral variance across tests, we employed dimension reduction via PCA of all baseline and end-point readouts. This analysis revealed that the top two principal components, explaining 30.7% of total variance, were strongly associated with stress-related outcomes, including physiological, anxiety-like, antidepressant-predictive, and cognitive deficits (based on parameter loadings). We then performed c-Fos mapping to characterize the functional consequences of chronic stress on the acute stress-mediated recruitment of 26 brain regions previously associated with stress responses. This analysis revealed that neuronal activation by acute stress was similarly exacerbated by CRS or UCMS in 4 out of 24 stress- and mood-related brain subregions investigated: the PFC A24b (ACC) and the ventral hippocampus CA1, CA3, and subiculum. Moreover, in regions dysregulated by chronic stress, levels of neuronal activation were positively correlated with stress-related principal components, suggesting that chronic stress-induced functional alterations in the recruitment of ACC or vHPC subregions by acute stress co-occur with or contribute to stress-related maladaptive behaviors.

As an advantage of our approach, we replicated findings of altered neuronal activation and maladaptive behaviors in parallel among two cohorts exposed to distinct stressor paradigms (i.e., CRS and UCMS), addressing key challenges in reliability and replicability associated with rodent chronic stress models and improving the likelihood that these results are meaningfully related to chronic stress-induced biological changes. Further, we combined a comprehensive phenotyping approach, using multiple tests assessing human symptom-like dimensions, with an unbiased dimension reduction technique to capture the full extent of behavioral variance attributable to chronic stress. This unbiased “bottom-up” PCA approach was well-suited for the current study to control for power bias introduced by an uneven number of tests loading onto individual parameters that may skew correlational analyses, e.g., relative to “top-down” methods for behavioral interpretation such as z-scoring^68^. We also included males and females in the current study based on evidence for a strong moderating role of sex in chronic stress processes^70,71^. However, biological outcomes were consistent across sexes and behavioral outcomes did not vastly differ in covariate analyses as, e.g., weight gain alterations were more severe in females week-to-week, but in males overall from week 0 to 8. Thus, we did not identify sex-specific effects, but cannot preclude this possibility due to the moderate sample size employed (6-7 males/females per group).

In this study, we confirmed that acute stress increases neuronal activation measured via c-Fos+ cell density across 18/24 stress- and mood-related brain regions examined, using an acute swim stressor. The rapid induction and short half-life of c-Fos make it an ideal marker to characterize functional changes in the acute stress response^47,65^. Numerous reports demonstrate that acute stress recruits regional activation via c-Fos expression patterns that overlap with our findings^72–75^. We found that prior chronic stress exposure exacerbated the effect of acute stress on neuronal activation of ACC and vHPC subregions. While this finding is interesting and relevant to reported dysregulations of these two brain regions in stress-related disorders^76–79^, past studies using prior exposure to chronic social or restraint stress reported contrasting attenuation of acute stress-induced c-Fos activation^44,48–50^. This discrepancy may be attributable to experimental design, as past studies used chronic stress protocols that were shorter (5-10 days), known to produce heterogeneous or resilient subgroups (e.g. social defeat)^80^, or that lacked validation with behavioral tests. Further, compared to our approach using a distinct acute stressor (i.e., cold swim vs. CRS/UCMS), past studies investigated response to the same stressor for acute and chronic phases, likely contributing to habituation^44,45^. For example, one study employing prior chronic social defeat stress before acute restraint stress revealed enhanced central amygdala activation, together with increased anxiety-like behavior^46^. However, we did not confirm this finding in the central amygdala, possibly due to lack of significant c-Fos induction by acute swim stress in this region, precluding insights about alterations in control vs. chronic stress-exposed mice. Consistent with our findings, another study found that c-Fos activation following chronic social defeat stress persisted at 24 hours in only 5/60 regions that were sensitive to activation by acute defeat, including the ACC and HPC^47^.

Although we confirmed enhanced activation of the ACC and vHPC, we did not fully replicate findings from studies using long-lasting IEG markers that demonstrate sustained activation by chronic stress in HPA-sensitive regions, such as the amygdala, PAG, and BNST^81–83^. These differences likely reflect our approach to measure functional response to acute stress vs. changes in baseline activity levels, but may have also been influenced by the type or intensity of stressors applied. Specifically, control vs. CRS/UCMS differences could have been masked by a “ceiling effect” on c-Fos activation due to the severity of stressor used, or else it may have been difficult to distinguish control mice due to repeated behavioral testing under stressful conditions. It is also important to mention that c-Fos expression is a proxy of neuronal activation and does not rule out the functional recruitment of brain regions through other mediators^97^. Finally, regional activity levels do not fully capture the complexity of circuit-level changes, wherein for instance, others showed that mPFC-vHPC activity becomes more tightly coupled in anxiogenic environments^98^.

Combining our findings in stress-naïve and chronic stress-exposed mice, we demonstrated that activation of the ACC and vHPC is recruited by acute stress, and that this effect is exacerbated by prior exposure to chronic stress. Parallel changes after chronic stress may reflect the fact that reciprocal connections between the ACC/vHPC (extending also to the amygdala, NAc, and BNST) exert important regulatory control of HPA-mediated stress responses^84^. Indeed, lesion studies targeting the PFC/ACC^85,86^ or vHPC/vSub^87^ demonstrated enhanced endocrine response to stress associated with greater c-Fos activation in GC-secreting structures. These findings are consistent with neuroimaging studies in healthy controls showing that smaller ACC volume was associated with enhanced HPA responses^88^. Similarly, HPC atrophy was associated with concomitant enhanced HPA responses and minor depressive symptoms in a study of AD patients^89^. Taken together, it seems that whereas acute recruitment of these brain regions is necessary for HPA regulatory feedback, chronic activation may negatively impact regulatory processes, leading to run-away downstream activity that support the transition from adaptive to pathological stress reactivity. Taken together, these findings provide strong justification for future studies to investigate functional changes using multiple indicators of neuronal activity (e.g., fMRI, short- and long-lasting IEG mapping, electrophysiology), and considering the dynamic interplay between markers of neuronal activation and endocrine stress response mediators both within and across functional brain circuits.

Although we did not assess specific cellular mediators, activity alterations are a likely consequence of deficits in excitatory and inhibitory neurons that are frequently observed after chronic stress. Indeed, human and animal studies focusing on the PFC/ACC and HPC revealed reductions in glutamatergic principal neuron morphology and synaptic connectivity that are reversed by rapid-acting antidepressant drugs, such as ketamine^6,116–118^. Similarly, altered structural and functional markers of inhibitory GABA neurons represent a pathological hallmark across disorders, are recapitulated by rodent chronic stress paradigms, and normalize following antidepressant treatment^33,119,120^. In contrast to our findings of enhanced excitation, multiple rodent studies reported that chronic stress increased GABAergic and decreased glutamatergic markers in the mPFC, potentially recapitulating prefrontal hypofunction in humans stress-related disorders (reviewed in ^90^). For example, 1-week repeated restraint or unpredictable stress reduced glutamatergic neurotransmission in mPFC pyramidal neurons in juvenile male rats^91^. Another study showed that 2-week exogenous corticosterone administration reduced vmPFC glutamate receptor subunits, which correlated with impaired fear extinction^92^. Finally, 2- or 4-week UCMS exposure increased GABAergic PV cell markers in the mPFC, but reduced local activation by novel environment stimuli only in female mice^93,94^. In light of these differences, it is important to distinguish that several past findings rely on stress paradigms shorter than the 2-week chronic stress minimum endorsed by a majority of researchers^51^. Further, a study using 9-week UCMS recapitulated human stress-related GABAergic deficits together with decreased inhibition of mPFC pyramidal neurons in susceptible rats^95^. Similarly, 3-weeks CRS exposure in rats exacerbated glutamatergic activation of HPC CA3 by an acute stressor^96^, consistent with our findings. Although these findings may seem discordant, they likely represent an imbalance attributable to the impact of chronic stress on both excitatory and inhibitory components. Although the precise mechanisms underlying these changes are unclear, one candidate is glutamatergic excitotoxicity^10,121^, supported by evidence that acute stress enhances extrasynaptic glutamate activity in the PFC and vHPC^5,122–125^, consistent with our c-Fos findings. This is supported by evidence that antidepressants and anxiolytics prevented the induction of stress-induced c-Fos activation in the ACC, HPC, CeA, PVN, and VTA^72,126–128^, supporting that dysregulation of the adaptive stress response precedes the structural and functional deficits that are targeted by pharmacological treatments.

In the current study, we identified altered ACC and vHPC activation together with physiological and anxiety-like deficits (CRS and UCMS) and impaired cognition (UCMS only). These findings are consistent with demonstrated roles for the ACC and vHPC in the regulation of mood and cognition, plus other functions that are disrupted in chronic stress-related disorders, including fear conditioning, reward processing, and motivation^76–79^. Indeed, postmortem studies of patients with MDD and BD demonstrate reduced gray matter volume and glial cell density in ACC A24b and adjacent subregions^99–101^. Elevated task-induced ACC activation has been identified in neuroimaging studies of patients with PTSD, anxiety disorders, and obsessive-compulsive disorder^102–105^. Although one meta-analysis revealed ACC hypoactivity in MDD (at baseline and during task-induction)^106^, others show that this was reversed when correcting for volumetric atrophy, revealing increased activity^37^. Reduced HPC volume, although moderate relative to ACC changes, was also consistently found in patients with MDD and PTSD^107–109^. In the HPC, findings are mixed. Postmortem studies in BD and MDD subjects demonstrated reduced synapse and gray matter volume in vCA1 and the subiculum, whereas imaging studies in live MDD patients reported volumetric changes in the HPC, but not when stratified to the anterior part (homologous to the vHPC in rodents)^110,111^. Functional changes also parallel those found in the ACC, showing hyperactivity during emotive processing in both regions, and associations between familial MDD risk and HPC hyperactivity that was reversed by antidepressants^112–115^.

Our study design revealed behavioral alterations consistent with past rodent chronic stress studies showing that >5-week CRS or UCMS exposure in rodents induces physiological alterations in weight gain and fur coat quality, anxiety-like behavior in the EPM, NSF, and PT, and cognitive impairments in working and short-term memory in the YM and NORT, respectively^19,20,129^. Comparisons of CRS and UCMS models often focus on differences in face and construct validity that parallel human disease experiences^58,129^. However, in line with recent Research Domain Criteria^130^, our approach aimed to focus on the convergent biological and behavioral alterations produced by both models to identify mechanisms connecting common disease risk factors (i.e., chronic stress) to maladaptive behaviors. Thus, we attribute the observed variability in behavioral alterations within and between models to well-documented challenges of using classical behavioral tests as measuring tools (see discussions on consistency, reliability, and variability in ^51,58,131^), rather than meaningful conclusions about the validity of UCMS vs. CRS. For example, we frequently observe YM deficits in CRS mice^57^, but did not here. To overcome these challenges, we implemented multiple end-point tests assessing top-down behavioral dimensions^131^ and used repeatable tests to (a) track the induction of deficits over time, and (b) provide baseline and endpoint readouts to measure the transition into maladaptive behavior (via PCA). Longitudinal tests revealed physiological and anxiety-like deficits persisting after 1 to 5 weeks of stress, supporting the utility of repeatable behavioral tests such as the PhenoTyper test^56,58,132^. Summary of trans-dimensional baseline and end-point behavioral alterations was performed using dimension reduction via principal component analysis based on past approaches^56,58^. We found that baseline parameters (e.g., week 0 or pre-trial readouts) had weak or negative loadings vs. the strong positive loadings of evaluated parameters (e.g., week 6-8 or trial readouts), indicating that these variables captured variance associated with maladaptive behavioral states. These PCs were positively correlated with c-Fos activation in regions altered by chronic stress, suggesting that dysregulation of the stress response in the ACC and vHPC co-occurs with or contributes to stress-related behaviors.

In summary, our data contribute evidence supporting a potential association between chronic stress, altered ACC and vHPC regional activation by acute stress, and maladaptive behaviors relevant to stress-related disorders. Although, the correlational design of the current study does not support causal inferences, future studies should investigate the modulation of excitatory and inhibitory components within these target regions to improve understanding of how chronic stress leads to pathological changes and to identify opportunities for treatment development.

## Acknowledgements & Funding Sources

CF, MB, and ES designed the study and wrote the manuscript. CF performed all behavior and IHC experiments with help from TP and MB. KM assisted with performing stressors. CF and TP were supported by CAMH Discovery Fund fellowships. CF also received an Ontario Graduate Scholarship during the studies. MB is supported by a NARSAD young investigator award from the Brain & Behavior Research Foundation (#24034) and the CAMH Discovery Seed Fund and the Canadian Institutes of Health Research (PJT-165852). ES was supported by the Brain & Behavior Research Foundation (#25637) and Canadian Institutes of Health Research (PJT-153175). The project was also supported by the Campbell Family Mental Health Research Institute.

## References

1. McEwen BS. Mood disorders and allostatic load. Biological Psychiatry 2003; 54: 200–207.

2. Ulrich-Lai YM, Herman JP. Neural regulation of endocrine and autonomic stress responses. Nature Reviews Neuroscience 2009; 10: 397–409.

3. Yuen EY, Liu W, Karatsoreos IN, et al. Acute stress enhances glutamatergic transmission in prefrontal cortex and facilitates working memory. Proc Natl Acad Sci US A 2009; 106: 14075–14079.

4. Moghaddam B. Stress Preferentially Increases Extraneuronal Levels of Excitatory Amino Acids in the Prefrontal Cortex: Comparison to Hippocampus and Basal Ganglia. J Neurochem 1993; 60: 1650–1657.

5. Lowy MT, Wittenberg L, Yamamoto BK. Effect of Acute Stress on Hippocampal Glutamate Levels and Spectrin Proteolysis in Young and Aged Rats. J Neurochem 1995; 65: 268–274.

6. Popoli M, Yan Z, McEwen BS, et al. The stressed synapse: the impact of stress and glucocorticoids on glutamate transmission. Nat Rev Neurosci 2012; 13: 22–37.

7. Duvarci S, Paré D. Glucocorticoids enhance the excitability of principal basolateral amygdala neurons. J Neurosci 2007; 27: 4482–4491.

8. Wang G-Y, Zhu Z-M, Cui S, et al. Glucocorticoid Induces Incoordination between Glutamatergic and GABAergic Neurons in the Amygdala. PLoS One 2016; 11: e0166535.

9. Di S, Maxson MM, Franco A, et al. Glucocorticoids regulate glutamate and GABA synapse-specific retrograde transmission via divergent nongenomic signaling pathways. J Neurosci 2009; 29:393–401.

10. Sousa N, Almeida OFX. Disconnection and reconnection: The morphological basis of (mal)adaptation to stress. Trends in Neurosciences 2012; 35: 742–751.

11. Shirazi SN, Friedman AR, Kaufer D, et al. Glucocorticoids and the brain: Neural mechanisms regulating the stress response. Adv Exp Med Biol 2015; 872: 235–252.

12. McEwen BS. Glucocorticoids, depression, and mood disorders: Structural remodeling in the brain. Metabolism 2005; 54: 20–23.

13. McEwen BS. Protection and damage from acute and chronic stress: Allostasis and allostatic overload and relevance to the pathophysiology of psychiatric disorders. New York Academy of Sciences.

14. Vyas S, Rodrigues AJ, Silva JM, et al. Chronic stress and glucocorticoids: From neuronal plasticity to neurodegeneration. Neural Plasticity; 2016. Epub ahead of print 2016. DOI: 10.1155/2016/6391686.

15. Schwabe L, Dickinson A, Wolf OT. Stress, habits, and drug addiction: a psychoneuroendocrinological perspective. Exp Clin Psychopharmacol 2011; 19: 53–63.

16. Girotti M, Adler SM, Bulin SE, et al. Prefrontal cortex executive processes affected by stress in health and disease. Progress in Neuro-Psychopharmacology and Biological Psychiatry 2018; 85: 161–179.

17. Shin LM, Liberzon I. The neurocircuitry of fear, stress, and anxiety disorders. Neuropsychopharmacology 2010; 35: 169–191.

18. Hill MN, Hellemans KGC, Verma P, et al. Neurobiology of chronic mild stress: parallels to major depression. Neurosci Biobehav Rev 2012; 36: 2085–117.

19. Mineur YS, Belzung C, Crusio WE. Effects of unpredictable chronic mild stress on anxiety and depression-like behavior in mice. Behav Brain Res 2006; 175: 43–50.

20. Chiba S, Numakawa T, Ninomiya M, et al. Chronic restraint stress causes anxiety- and depression-like behaviors, downregulates glucocorticoid receptor expression, and attenuates glutamate release induced by brain-derived neurotrophic factor in the prefrontal cortex. Prog Neuropsychopharmacol Biol Psychiatry 2012; 39: 112–9.

21. Nollet M, Le Guisquet A-M, Belzung C. Models of depression: unpredictable chronic mild stress in mice. Curr Protoc Pharmacol 2013; Chapter 5: Unit 5.65.

22. Cook SC, Wellman CL. Chronic stress alters dendritic morphology in rat medial prefrontal cortex. J Neurobiol 2004; 60: 236–248.

23. Wellman CL. Dendritic reorganization in pyramidal neurons in medial prefrontal cortex after chronic corticosterone administration. J Neurobiol 2001; 49: 245–253.

24. Radley JJ, Sisti HM, Hao J, et al. Chronic behavioral stress induces apical dendritic reorganization in pyramidal neurons of the medial prefrontal cortex. Neuroscience 2004; 125: 1–6.

25. Banasr M, Duman RS. Glial Loss in the Prefrontal Cortex Is Sufficient to Induce Depressive-like Behaviors. Biol Psychiatry 2008; 64: 863–870.

26. Zhang H, Zhao Y, Wang Z. Chronic corticosterone exposure reduces hippocampal astrocyte structural plasticity and induces hippocampal atrophy in mice. Neurosci Lett 2015; 592: 76–81.

27. Magarin os AM, McEwen BS. Stress-induced atrophy of apical dendrites of hippocampal CA3c neurons: Comparison of stressors. Neuroscience 1995; 69: 83–88.

28. Vyas A, Mitra R, Shankaranarayana Rao BS, et al. Chronic stress induces contrasting patterns of dendritic remodeling in hippocampal and amygdaloid neurons. 2002; 22: 6810–6818.

29. Vyas A, Bernal S, Chattarji S. Effects of chronic stress on dendritic arborization in the central and extended amygdala. Brain Res 2003; 965: 290–294.

30. Kang HJ, Voleti B, Hajszan T, et al. Decreased expression of synapse-related genes and loss of synapses in major depressive disorder. Nat Med 2012; 18: 1413–7.

31. Sheline YI, Wang PW, Gado MH, et al. Hippocampal atrophy in recurrent major depression. Proc Natl Acad Sci U S A 1996; 93: 3908–3913.

32. Khundakar A, Morris C, Oakley A, et al. Morphometry analysis of neuronal and glial cell pathology in the dorsolateral prefrontal cortex in late-life depression. Br J Psychiatry 2009; 195: 163–169.

33. Fee C, Banasr M, Sibille E. Somatostatin-Positive Gamma-Aminobutyric Acid Interneuron Deficits in Depression: Cortical Microcircuit and Therapeutic Perspectives. Biol Psychiatry 2017; 82: 549–559.

34. Sullivan RM, Gratton A. Prefrontal cortical regulation of hypothalamic-pituitary-adrenal function in the rat and implications for psychopathology: Side matters. Psychoneuroendocrinology 2002; 27:99–114.

35. Sapolsky RM. A mechanism for glucocorticoid toxicity in the hippocampus: Increased neuronal vulnerability to metabolic insults. J Neurosci 1985; 5: 1228–1232.

36. McEwen BS, Gianaros PJ. Stress- and Allostasis-Induced Brain Plasticity. Annu Rev Med 2011; 62:431–445.

37. Drevets WC, Price JL, Furey ML. Brain structural and functional abnormalities in mood disorders: implications for neurocircuitry models of depression. Brain Struct Funct 2008; 213: 93–118.

38. Pitman RK, Rasmusson AM, Koenen KC, et al. Biological studies of post-traumatic stress disorder. Nature Reviews Neuroscience 2012; 13: 769–787.

39. Henckens MJAG, van der Marel K, van der Toorn A, et al. Stress-induced alterations in large-scale functional networks of the rodent brain. Neuroimage 2015; 105: 312–22.

40. Jacob Y, Morris LS, Huang KH, et al. Neural correlates of rumination in major depressive disorder: A brain network analysis. NeuroImage Clin; 25. Epub ahead of print 1 January 2020. DOI: 10.1016/j.nicl.2019.102142.

41. Nathan DE, Bellgowan JAF, French LM, et al. Assessing the Impact of Post-Traumatic Stress Symptoms on the Resting-State Default Mode Network in a Military Chronic Mild Traumatic Brain Injury Sample. Brain Connect 2017; 7: 236–249.

42. Lin H, Cai X, Zhang D, et al. Functional connectivity markers of depression in advanced Parkinson’s disease. NeuroImage Clin; 25. Epub ahead of print 1 January 2020. DOI: 10.1016/j.nicl.2019.102130.

43. Zhang R, Volkow ND. Brain default-mode network dysfunction in addiction. NeuroImage 2019; 200:313–331.

44. Mohammad G, Chowdhury I, Fujioka T, et al. Induction and adaptation of Fos expression in the rat brain by two types of acute restraint stress. Brain Res Bull 2000; 52: 171–182.

45. Figueiredo HF, Bruestle A, Bodie B, et al. The medial prefrontal cortex differentially regulates stress-induced c-fos expression in the forebrain depending on type of stressor. Eur J Neurosci 2003;18:2357–2364.

46. Chung KKK, Martinez M, Herbert J. c-fos expression, behavioural, endocrine and autonomic responses to acute social stress in male rats after chronic restraint: modulation by serotonin. Neuroscience 1999; 95: 453–463.

47. Matsuda S, Peng H, Yoshimura H, et al. Persistent c-fos expression in the brains of mice with chronic social stress. Neurosci Res 1996; 26: 157–170.

48. Umemoto S, Noguchi K, Kawai Y, et al. Repeated stress reduces the subsequent stress-induced expression of Fos in rat brain. Neurosci Lett 1994; 167: 101–104.

49. Stamp JA, Herbert J. Multiple immediate-early gene expression during physiological and endocrine adaptation to repeated stress. Neuroscience 1999; 94: 1313–1322.

50. Melia KR, Ryabinin AE, Schroeder R, et al. Induction and habituation of immediate early gene expression in rat brain by acute and repeated restraint stress. J Neurosci 1994; 14: 5929–5938.

51. Willner P. Reliability of the chronic mild stress model of depression: A user survey. Neurobiol Stress 2017; 6: 68–77.

52. Bagot RC, Parise EM, Peña CJ, et al. Ventral hippocampal afferents to the nucleus accumbens regulate susceptibility to depression. Nat Commun; 6. Epub ahead of print 8 May 2015. DOI: 10.1038/ncomms8062.

53. Russo SJ, Nestler EJ. The brain reward circuitry in mood disorders. Nature Reviews Neuroscience 2013; 14:609–625.

54. Tynan RJ, Naicker S, Hinwood M, et al. Chronic stress alters the density and morphology of microglia in a subset of stress-responsive brain regions. Brain Behav Immun 2010; 24: 1058–68.

55. Kim Y, Perova Z, Mirrione MM, et al. Whole-brain mapping of neuronal activity in the learned helplessness model of depression. Front Neural Circuits; 10. Epub ahead of print 3 February 2016. DOI: 10.3389/fncir.2016.00003.

56. Nikolova YS, Misquitta KA, Rocco BR, et al. Shifting priorities: highly conserved behavioral and brain network adaptations to chronic stress across species. Transl Psychiatry 2018; 8: 26.

57. Prevot TD, Li G, Vidojevic A, et al. Novel Benzodiazepine-Like Ligands with Various Anxiolytic, Antidepressant, or Pro-Cognitive Profiles. Mol Neuropsychiatry 2019; 5: 84–97.

58. Prevot TD, Misquitta KA, Fee C, et al. Residual avoidance: A new, consistent and repeatable readout of chronic stress-induced conflict anxiety reversible by antidepressant treatment. Neuropharmacology 2019; 153: 98–110.

59. Isingrini E, Camus V, Le Guisquet AM, et al. Association between repeated unpredictable chronic mild stress (UCMS) procedures with a high fat diet: A model of fluoxetine resistance in mice. PLoS One; 5. Epub ahead of print 2010. DOI: 10.1371/journal.pone.0010404.

60. Samuels BA, Hen R. Novelty-suppressed feeding in the mouse. Neuromethods 2011; 63: 107–121.

61. Dulawa SC, Hen R. Recent advances in animal models of chronic antidepressant effects: The novelty-induced hypophagia test. Neuroscience and Biobehavioral Reviews 2005; 29: 771–783.

62. Slattery DA, Cryan JF. Using the rat forced swim test to assess antidepressant-like activity in rodents. Nature Protocols 2012; 7: 1009–1014.

63. Olton DS. Mazes, maps, and memory. Am Psychol 1979; 34: 583–596.

64. Antunes M, Biala G. The novel object recognition memory: Neurobiology, test procedure, and its modifications. Cognitive Processing 2012; 13: 93–110.

65. Kovács KJ. c-Fos as a transcription factor: A stressful (re)view from a functional map. Neurochemistry International 1998; 33: 287–297.

66. Franklin K, Paxinos G. The mouse brain in stereotaxic coordinates.

67. Wheeler AL, Teixeira CM, Wang AH, et al. Identification of a Functional Connectome for Long Term Fear Memory in Mice. PLoS Comput Biol 2013; 9: e1002853.

68. Guilloux J-P, Seney M, Edgar N, et al. Integrated behavioral z-scoring increases the sensitivity and reliability of behavioral phenotyping in mice: relevance to emotionality and sex. 2011; 197: 21–31.

69. Carola V, D’Olimpio F, Brunamonti E, et al. Evaluation of the elevated plus-maze and open-field tests for the assessment of anxiety-related behaviour in inbred mice. Behav Brain Res 2002; 134: 49–57.

70. Cahill L. Why sex matters for neuroscience. Nature Reviews Neuroscience 2006; 7: 477–484.

71. Garrett JE, Wellman CL. Chronic stress effects on dendritic morphology in medial prefrontal cortex: sex differences and estrogen dependence. Neuroscience 2009; 162: 195–207.

72. Silva M, Aguiar DC, Diniz CRA, et al. Neuronal NOS inhibitor and conventional antidepressant drugs attenuate stress-induced fos expression in overlapping brain regions. Cell Mol Neurobiol 2012; 32:443–53.

73. Senba E, Ueyama T. Stress-induced expression of immediate early genes in the brain and peripheral organs of the rat. Neurosci Res 1997; 29: 183–207.

74. Senba E, Umemoto S, Kawai Y, et al. Differential expression of fos family and jun family mRNAs in the rat hypothalamo-pituitary-adrenal axis after immobilization stress. Brain Res Mol Brain Res 1994; 24: 283–94.

75. Bubser M, Deutch AY. Stress induces Fos expression in neurons of the thalamic paraventricular nucleus that innervate limbic forebrain sites. Synapse 1999; 32: 13–22.

76. Etkin A, Egner T, Kalisch R. Emotional processing in anterior cingulate and medial prefrontal cortex. Trends in Cognitive Sciences 2011; 15: 85–93.

77. Quirk GJ, Beer JS. Prefrontal involvement in the regulation of emotion: convergence of rat and human studies. Current Opinion in Neurobiology 2006; 16: 723–727.

78. Tannenholz L, Jimenez JC, Kheirbek MA. Local and regional heterogeneity underlying hippocampal modulation of cognition and mood. Frontiers in Behavioral Neuroscience; 8. Epub ahead of print 6 May 2014. DOI: 10.3389/fnbeh.2014.00147.

79. Phan KL, Wager T, Taylor SF, et al. Functional neuroanatomy of emotion: A meta-analysis of emotion activation studies in PET and fMRI. NeuroImage 2002; 16: 331–348.

80. Krishnan V, Han MH, Graham DL, et al. Molecular Adaptations Underlying Susceptibility and Resistance to Social Defeat in Brain Reward Regions. Cell 2007; 131: 391–404.

81. Laine MA, Sokolowska E, Dudek M, et al. Brain activation induced by chronic psychosocial stress in mice. Sci Rep; 7. Epub ahead of print 1 December 2017. DOI: 10.1038/s41598-017-15422-5.

82. Perrotti LI, Hadeishi Y, Ulery PG, et al. Induction of ΔFosB in reward-related brain structures after chronic stress. J Neurosci 2004; 24: 10594–10602.

83. Flak JN, Solomon MB, Jankord R, et al. Identification of chronic stress-activated regions reveals a potential recruited circuit in rat brain. Eur J Neurosci 2012; 36: 2547–2555.

84. Bannerman DM, Rawlins JNP, McHugh SB, et al. Regional dissociations within the hippocampus - Memory and anxiety. Neuroscience and Biobehavioral Reviews 2004; 28: 273–283.

85. Diorio D, Viau V, Meaney MJ. The role of the medial prefrontal cortex (cingulate gyrus) in the regulation of hypothalamic-pituitary-adrenal responses to stress. J Neurosci 1993; 13: 3839–3847.

86. Radley JJ, Arias CM, Sawchenko PE. Regional differentiation of the medial prefrontal cortex in regulating adaptive responses to acute emotional stress. J Neurosci 2006; 26: 12967–12976.

87. Jankord R, Herman JP. Limbic regulation of hypothalamo-pituitary-adrenocortical function during acute and chronic stress. In: Annals of the New York Academy of Sciences. Blackwell Publishing Inc., 2008, pp. 64–73.

88. MacLullich AMJ, Ferguson KJ, Wardlaw JM, et al. Smaller left anterior cingulate cortex volumes are associated with impaired hypothalamic-pituitary-adrenal axis regulation in healthy elderly men. J Clin Endocrinol Metab 2006; 91: 1591–1594.

89. O’Brien JT, Ames D, Schweitzer I, et al. Clinical and magnetic resonance imaging correlates of hypothalamic-pituitary-adrenal axis function in depression and Alzheimer’s disease. Br J Psychiatry 1996; 168: 679–687.

90. McKlveen JM, Moloney RD, Scheimann JR, et al. “Braking” the Prefrontal Cortex: The Role of Glucocorticoids and Interneurons in Stress Adaptation and Pathology. Biological Psychiatry 2019; 86: 669–681.

91. Yuen EY, Wei J, Liu W, et al. Repeated Stress Causes Cognitive Impairment by Suppressing Glutamate Receptor Expression and Function in Prefrontal Cortex. Neuron 2012; 73: 962–977.

92. Gourley SL, Kedves AT, Olausson P, et al. A history of corticosterone exposure regulates fear extinction and cortical NR2B, GluR2/3, and BDNF. Neuropsychopharmacology 2009; 34: 707–716.

93. Shepard R, Page CE, Coutellier L. Sensitivity of the prefrontal GABAergic system to chronic stress in male and female mice: Relevance for sex differences in stress-related disorders. Neuroscience 2016; 332: 1–12.

94. Shepard R, Coutellier L. Changes in the Prefrontal Glutamatergic and Parvalbumin Systems of Mice Exposed to Unpredictable Chronic Stress. Mol Neurobiol 2018; 55: 2591–2602.

95. Czéh B, Vardya I, Varga Z, et al. Long-term stress disrupts the structural and functional integrity of GABAergic neuronal networks in the medial prefrontal cortex of rats. Front Cell Neurosci; 12. Epub ahead of print 20 June 2018. DOI: 10.3389/fncel.2018.00148.

96. Yamamoto BK, Reagan LP. The glutamatergic system in neuronal plasticity and vulnerability in mood disorders. Neuropsychiatr Dis Treat 2006; 2: 7–14.

97. Ericsson A, Kovács KJ, Sawchenko PE. A functional anatomical analysis of central pathways subserving the effects of interleukin-1 on stress-related neuroendocrine neurons. J Neurosci 1994; 14:897–913.

98. Adhikari A, Topiwala MA, Gordon JA. Synchronized Activity between the Ventral Hippocampus and the Medial Prefrontal Cortex during Anxiety. Neuron 2010; 65: 257–269.

99. Öngür D, Drevets WC, Price JL. Glial reduction in the subgenual prefrontal cortex in mood disorders. Proc Natl Acad Sci U S A 1998; 95: 13290–13295.

100. Drevets WC, Savitz J, Trimble M. The subgenual anterior cingulate cortex in mood disorders. CNS Spectr 2008; 13: 663–681.

101. Cotter D, Mackay D, Landau S, et al. Reduced glial cell density and neuronal size in the anterior cingulate cortex in major depressive disorder. Arch Gen Psychiatry 2001; 58: 545–553.

102. Fitzgerald KD, Welsh RC, Gehring WJ, et al. Error-related hyperactivity of the anterior cingulate cortex in obsessive-compulsive disorder. Biol Psychiatry 2005; 57: 287–294.

103. McClure EB, Monk CS, Nelson EE, et al. Abnormal attention modulation of fear circuit function in pediatric generalized anxiety disorder. Arch Gen Psychiatry 2007; 64: 97–106.

104. Nitschke JB, Sarinopoulos I, Oathes DJ, et al. Anticipatory activation in the Amygdala and Anterior Cingulate in generalized anxiety disorder and prediction of reatment response. Am J Psychiatry 2009; 166: 302–310.

105. Etkin A, Wager TD. Functional neuroimaging of anxiety: A meta-ana lysis of emotional processing in PTSD, social anxiety disorder, and specific phobia. American Journal of Psychiatry 2007;164: 1476–1488.

106. Fitzgerald PB, Laird AR, Maller J, et al. A meta-analytic study of changes in brain activation in depression. Hum Brain Mapp 2008; 29: 683–695.

107. Koolschijn PCMP, van Haren NEM, Lensvelt-Mulders GJLM, et al. Brain volume abnormalities in major depressive disorder: A meta-analysis of magnetic resonance imaging studies. Hum Brain Mapp 2009; 30: 3719–3735.

108. Stratmann M, Konrad C, Kugel H, et al. Insular and Hippocampal Gray Matter Volume Reductions in Patients with Major Depressive Disorder. PLoS One 2014; 9: e102692.

109. Henigsberg N, Kalember P, Petrović ZK, et al. Neuroimaging research in posttraumatic stress disorder – Focus on amygdala, hippocampus and prefrontal cortex. Progress in Neuro-Psychopharmacology and Biological Psychiatry 2019; 90: 37–42.

110. Neumeister A, Wood S, Bonne O, et al. Reduced hippocampal volume in unmedicated, remitted patients with major depression versus control subjects. Biol Psychiatry 2005; 57: 935–937.

111. Bonne O, Vythilingam M, Inagaki M, et al. Reduced posterior hippocampal volume in posttraumatic stress disorder. J Clin Psychiatry 2008; 69: 1087–1091.

112. Jaworska N, Yang X-R, Knott V, et al. A review of fMRI studies during visual emotive processing in major depressive disorder. 16. Epub ahead of print 3 October 2015. DOI: 10.3109/15622975.2014.885659.

113. Mayberg HS, Brannan SK, Tekell JL, et al. Regional metabolic effects of fluoxetine in major depression: Serial changes and relationship to clinical response. In: Biological Psychiatry. 2000, pp. 830–843.

114. Kennedy SH, Evans KR, Krüger S, et al. Changes in regional brain glucose metabolism measured with positron emission tomography after paroxetine treatment of major depression. Am J Psychiatry 2001; 158: 899–905.

115. Mannie ZN, Filippini N, Williams C, et al. Structural and functional imaging of the hippocampus in young people at familial risk of depression. Psychol Med 2014; 44: 2939–2948.

116. Li N, Liu R-J, Dwyer JM, et al. Glutamate N-methyl-D-aspartate Receptor Antagonists Rapidly Reverse Behavioral and Synaptic Deficits Caused by Chronic Stress Exposure. Biol Psychiatry 2011; 69: 754–761.

117. McEwen BS, Bowles NP, Gray JD, et al. Mechanisms of stress in the brain. Nat Neurosci 2015; 18:1353–1363.

118. Zhao J, Verwer RWH, van Wamelen DJ, et al. Prefrontal changes in the glutamate-glutamine cycle and neuronal/glial glutamate transporters in depression with and without suicide. J Psychiatr Res 2016; 82:8–15.

119. Banasr M, Lepack A, Fee C, et al. Characterization of GABAergic Marker Expression in the Chronic Unpredictable Stress Model of Depression. Chronic Stress 2017; 1: 247054701772045.

120. Ghosal S, Duman CH, Liu R-J, et al. Ketamine rapidly reverses stress-induced impairments in GABAergic transmission in the prefrontal cortex in male rodents. Neurobiol Dis 2020; 134: 104669.

121. Duman RS, Aghajanian GK, Sanacora G, et al. Synaptic plasticity and depression: new insights from stress and rapid-acting antidepressants. Nat Med 2016; 22: 238–249.

122. Yuen EY, Liu W, Karatsoreos IN, et al. Mechanisms for acute stress-induced enhancement of glutamatergic transmission and working memory. Mol Psychiatry 2011; 16: 156–170.

123. Treccani G, Musazzi L, Perego C, et al. Stress and corticosterone increase the readily releasable pool of glutamate vesicles in synaptic terminals of prefrontal and frontal cortex. Mol Psychiatry 2014; 19:433–443.

124. Maggio N, Segal M. Differential modulation of long-term depression by acute stress in the rat dorsal and ventral hippocampus. J Neurosci 2009; 29: 8633–8638.

125. Venero C, Borrell J. Rapid glucocorticoid effects on excitatory amino acid levels in the hippocampus: a microdialysis study in freely moving rats. Eur J Neurosci 1999; 11: 2465–2473.

126. Morrow BA, Elsworth JD, Lee EJK, et al. Divergent effects of putative anxiolytics on stress-induced Fos expression in the mesoprefrontal system of the rat. Synapse 2000; 36: 143–154.

127. Beck CHM, Fibiger HC. Chronic desipramine alters stress-induced behaviors and regional expression of the immediate early gene, c-fos. Pharmacol Biochem Behav 1995; 51: 331–338.

128. Lino-de-Oliveira C, Sales AJ, Aparecida Del Bel E, et al. Effects of acute and chronic fluoxetine treatments on restraint stress-induced Fos expression. Brain Res Bull 2001; 55: 747–754.

129. Willner P. The chronic mild stress (CMS) model of depression: History, evaluation and usage. Neurobiology of Stress 2017; 6: 78–93.

130. Anderzhanova E, Kirmeier T, Wotjak CT. Animal models in psychiatric research: The RDoC system as a new framework for endophenotype-oriented translational neuroscience. Neurobiology of Stress 2017; 7: 47–56.

131. Ramos A. Animal models of anxiety: do I need multiple tests? Trends Pharmacol Sci 2008; 29: 493–498.

132. Maluach AM, Misquitta KA, Prevot TD, et al. Increased Neuronal DNA/RNA Oxidation in the Frontal Cortex of Mice Subjected to Unpredictable Chronic Mild Stress. Chronic Stress 2017; 1: 247054701772474.

