## Supplementary Table and Figures for "Chronic stress exacerbates acute stress-induced neuronal activation in the anterior cingulate cortex and ventral hippocampus that correlates with behavioral deficits in mice"

**Running title**: Dysregulated neuronal activation & behavioral deficits

**Suppl. Table 1.** Significant influence of sex on alterations in weight gain, coat state, PhenoTyper Test (PT) shelter zone area under the curve (AUC), and novelty-induced hypophagia (NIH) latency to drink novel assessed in control, CRS, and UCMS mice.

|  | **Control** | | | | **CRS** | | | | **UCMS** | | | | **Males Only** | | **Females Only** | |
| --- | --- | --- | --- | --- | --- | --- | --- | --- | --- | --- | --- | --- | --- | --- | --- | --- |
|  | Male | | Female | | Male | | Female | | Male | | Female | |  |  |  |  |
|  | Mean | ±SEM | Mean | ±SEM | Mean | ±SEM | Mean | ±SEM | Mean | ±SEM | Mean | ±SEM | Mean | ±SEM | Mean | ±SEM |
| **Weight Gain (% change)** | | | | | | | | | | | | | | | | |
| Week 1 | 2.06 | 0.49 | 5.61 | 1.18 | -2.82 | 1.44 | 2.49 | 0.67 | 1.47 | 0.70 | 6.93 | 0.74 | 0.33 | 0.71 | 4.91 | 0.66 |
| Week 2 | 1.47 | 0.64 | 4.59 | 0.77 | 2.76 | 1.30 | 3.53 | 1.20 | -1.99 | 1.50 | 2.04 | 0.82 | 0.79 | 0.78 | 3.46 | 0.58 |
| Week 3 | 0.17 | 0.41 | 1.07 | 0.55 | 0.28 | 0.40 | 0.85 | 0.79 | 0.63 | 0.28 | 0.25 | 0.38 | 0.35 | 0.21 | 0.75 | 0.35 |
| Week 4 | -0.10 | 0.49 | -1.94 | 0.94 | -4.18 | 0.73 | -3.30 | 0.90 | 0.64 | 1.80 | -1.38 | 0.76 | -1.15 | 0.78 | -2.25 | 0.52 |
| Week 5 | 0.47 | 0.72 | 0.94 | 1.43 | -2.39 | 0.71 | -1.49 | 0.56 | -0.37 | 0.62 | 0.16 | 0.61 | -0.70 | 0.47 | -0.14 | 0.59 |
| Week 0-8 Total | 11.01 | 1.88 | 18.59 | 0.38 | 1.44 | 1.18 | 10.51 | 1.11 | 2.58 | 1.06 | 17.28 | 1.37 | 5.33 | 1.31 | 15.37 | 0.99 |
| **Coat State (A.U.)** | | | | | | | | | | | | | | | | |
| Week 0 | 0.14 | 0.09 | 0.00 | 0.00 | 0.42 | 0.15 | 0.00 | 0.00 | 0.25 | 0.17 | 0.00 | 0.00 | 0.26 | 0.08 | 0.00 | 0.00 |
| Week 1 | 0.14 | 0.09 | 0.00 | 0.00 | 3.17 | 0.36 | 0.79 | 0.29 | 2.58 | 0.40 | 0.42 | 0.08 | 1.87 | 0.35 | 0.40 | 0.12 |
| Week 2 | 1.21 | 0.24 | 0.00 | 0.00 | 4.67 | 0.25 | 1.36 | 0.18 | 3.42 | 0.35 | 0.75 | 0.21 | 3.00 | 0.37 | 0.70 | 0.16 |
| Week 3 | 1.07 | 0.17 | 0.00 | 0.00 | 4.67 | 0.17 | 1.64 | 0.18 | 3.83 | 0.51 | 1.08 | 0.15 | 3.08 | 0.41 | 0.90 | 0.18 |
| Week 4 | 1.86 | 0.30 | 0.07 | 0.07 | 5.50 | 0.41 | 3.50 | 0.15 | 4.42 | 0.30 | 1.50 | 0.55 | 3.82 | 0.41 | 1.70 | 0.37 |
| Week 5 | 1.43 | 0.20 | 0.07 | 0.07 | 5.58 | 0.44 | 3.07 | 0.23 | 4.67 | 0.31 | 1.75 | 0.50 | 3.76 | 0.46 | 1.63 | 0.33 |
| Week 0-8 Total | 1.36 | 0.21 | 0.07 | 0.07 | 5.75 | 0.46 | 3.07 | 0.30 | 5.25 | 0.56 | 2.17 | 0.46 | 3.97 | 0.52 | 1.75 | 0.34 |
| **PT Shelter Zone Time AUC (s, 10^3)** | | | | | | | | | | | | | | | | |
|  | 15.31 | 0.73 | 12.99 | 1.23 | 18.76 | 1.23 | 18.20 | 0.37 | 20.85 | 0.66 | 17.50 | 0.94 | 18.15 | 0.73 | 16.17 | 0.73 |
| **NIH Latency to Drink Novel (s)** | | | | | | | | | | | | | | | | |
|  | 43.43 | 5.36 | 47.43 | 10.27 | 36.00 | 7.29 | 60.29 | 13.06 | 45.50 | 6.88 | 79.33 | 15.69 | 41.74 | 3.64 | 61.50 | 7.66 |

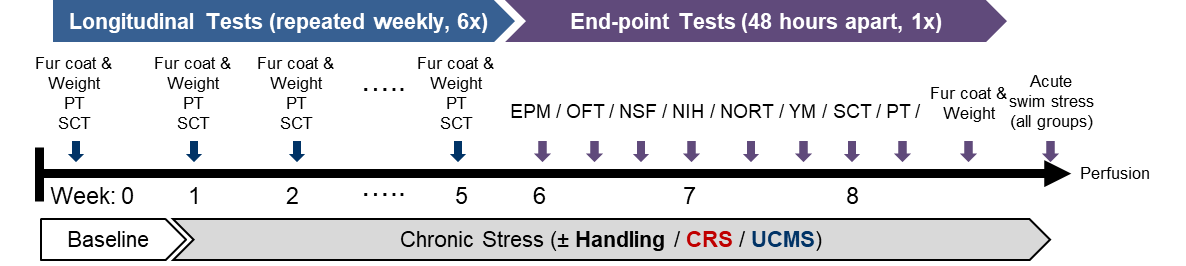

**Suppl. Fig. 1.** Study design for longitudinal and end-point behavioral analysis plus acute stressor challenge prior to brain perfusion. Longitudinal tests (i.e., repeatable tests not relying on novelty) confirmed the induction of chronic stress-related behavioral deficits from week 0 (baseline) to week 5 (the 5^th^ week of chronic stress). End-point tests characterized the full extent of depressive-like deficits from weeks 6-8 (after 5+ weeks chronic stress) based on an order that we find to be reliable to detect responses. PhenoTyper test (PT), Sucrose Consumption Test (SCT), Elevated Plus-maze (EPM), Open Field Test (OFT), Novelty-Suppressed Feeding (NSF), Novelty-Induced Hypophagia (NIH), Novel Object Recognition Test (NORT), Y-maze (YM), Chronic Restraint Stress (CRS), Unpredictable Chronic Mild Stress (UCMS).

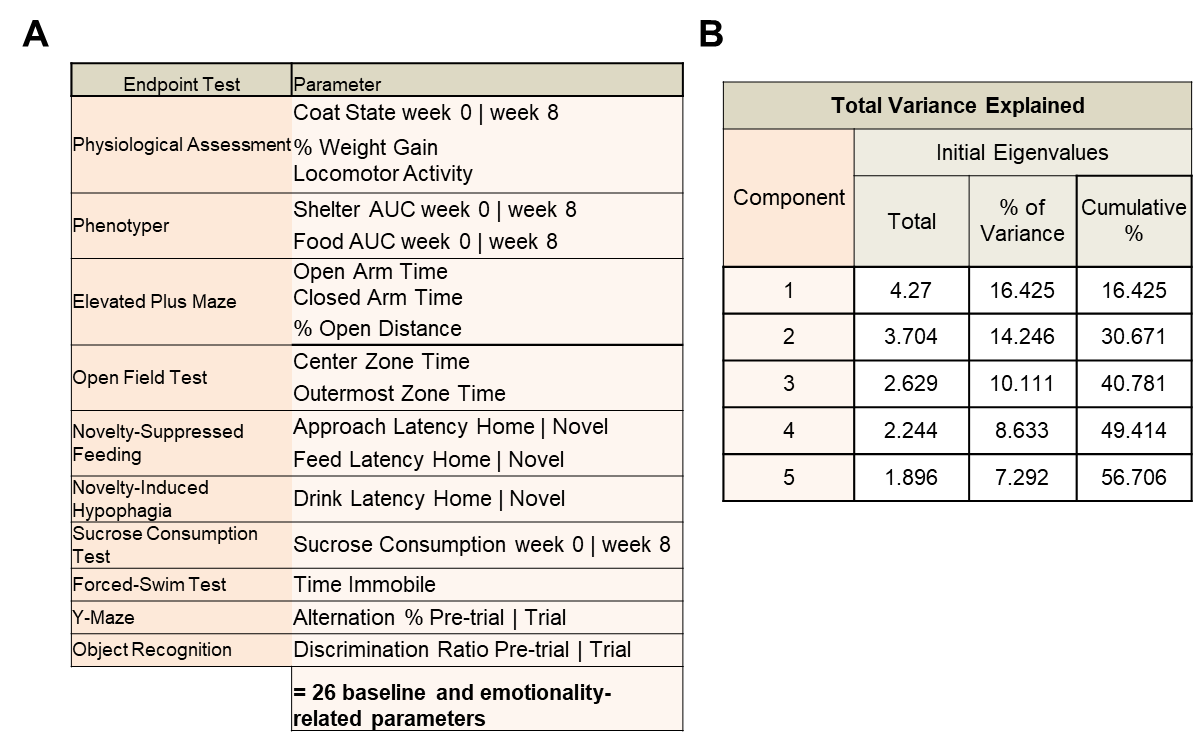

**Suppl. Fig. 2.** (A) list of behavioral parameters loaded into principal components analysis (PCA). Baseline parameters = week 0 or pre-trial readouts (e.g., latency to feed in a home cage test), end-point parameters = weeks 6-8 trial readouts (e.g., latency to feed in a novel cage test). (B) PCA initial eigenvalues representing the percentage variance captured for the top 5 components (>50% of total variance) in control, CRS, and UCMS mice.

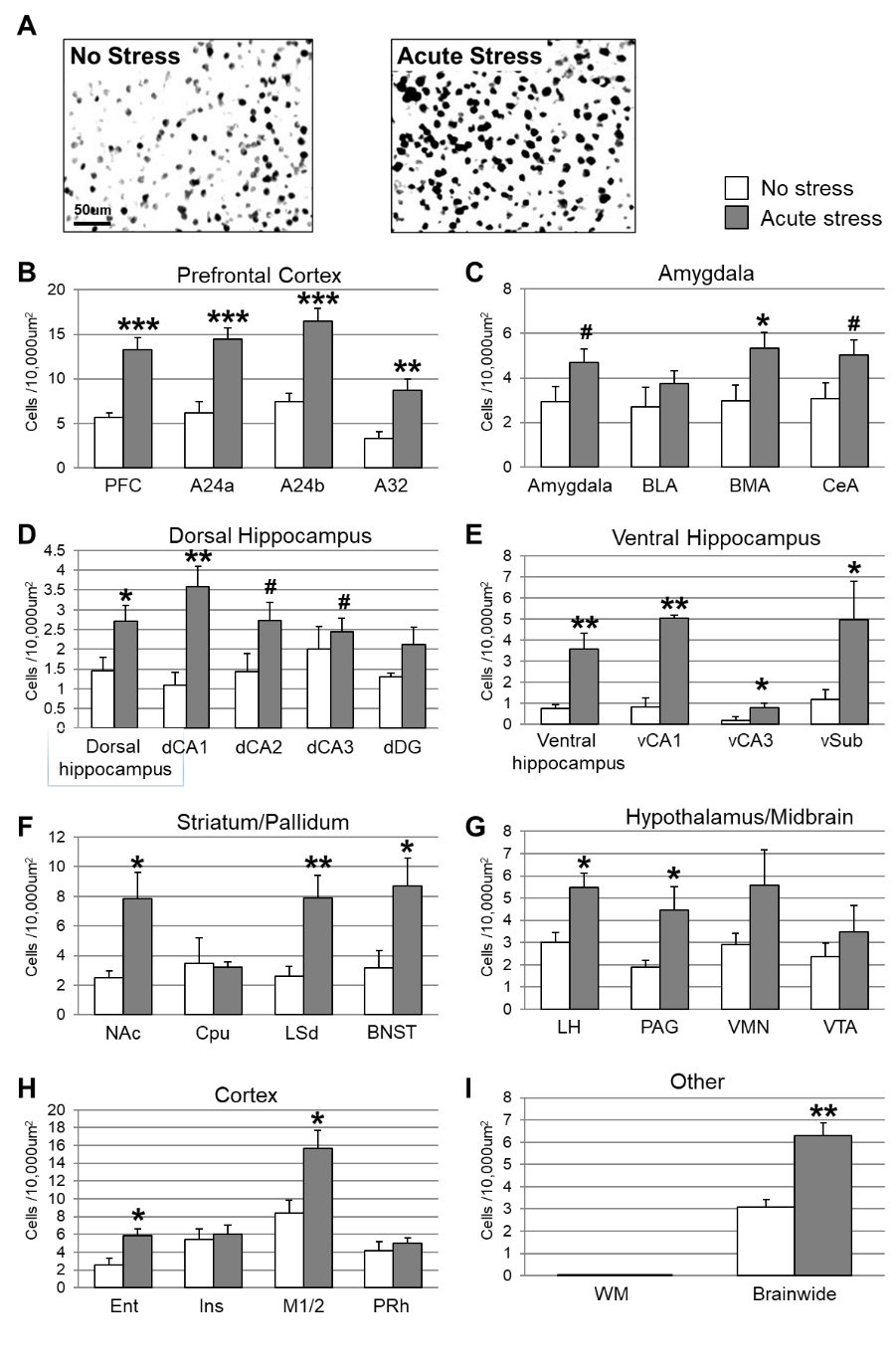

**Suppl. Fig. 3.** Neuronal activation quantified regionally via absolute density counts for c-Fos+ cells 90 min after exposure to no stress (white) or acute stress (*n*=6/group; 50% male/female). (A) Representative qualitative images of c-Fos staining in A24b. (B) Prefrontal cortex (PFC) as an average index of A24a, A24b, and A32. (C) Amygdala as an average index of basolateral (BLA), basomedial (BMA), and central (CeA) parts. (D) Dorsal hippocampus (dHPC) as an average index of CA1, CA2, CA3, and dentate gyrus (DG). (E) Ventral hippocampus (vHPC) as an average index of CA1, CA3, and subiculum (sub). (F) Striatum/pallidum groups comprising nucleus accumbens (NAc), caudate putamen (Cpu), lateral septum (LSd), and bed nucleus of stria terminalis (BNST). (G) Hypothalamus/midbrain groups comprising lateral hypothalamus (LH), periaqueductal gray (PAG), ventromedial nucleus (VMN), and ventral tegmental area (VTA). (H) Other cortical regions including entorhinal (Ent), insular (Ins), motor (M1/2), and perirhinal (PRh) cortex. (I) Control and overall regions: white matter (WM) and brain-wide (average of all regions). *** *p* < .001; ** *p* < .01; * *p* < .05 for indicated group vs. controls.
